# Emergence of a senescent and inflammatory pulmonary CD4^+^ T cell population prior to lung allograft failure

**DOI:** 10.1101/2023.12.19.572416

**Authors:** Sajad Moshkelgosha, Liran Levy, Shahideh Safavi, Sumiha Karunagaran, Gavin Wilson, Benjamin Renaud-Picard, Goodness Madu, Rashi Ramchandani, Jillian Oliver, Tatsuaki Watanabe, Ke Fan Bei, Betty Joe, Qixuan Li, Ella Huszti, May Cheung, David Hedley, Jonathan Yeung, Shaf Keshavjee, Tereza Martinu, Stephen Juvet

**Author notes:** Address of Corresponding Author: Stephen Juvet, MD, PhD Toronto General Hospital: University Health Network (UHN) 200 Elizabeth St., Rm 11 PMB 126 Toronto, Ontario, M5G 2C4 - ext: 8178. **<u>Disclosure</u>** Sajad Moshkelgosha, Shaf Keshavjee and Stephen Juvet have filed US patent US63/460,274 in relation to the work in this manuscript.

## Abstract

Lung transplantation is a life-saving therapy for end-stage pulmonary disease, but its long-term outlook is poor due to a high incidence of chronic lung allograft dysfunction (CLAD). CLAD results from alloimmune-mediated progressive fibrosis and culminates in death or the need for re-transplantation after a median of 6 years. Existing immunosuppression fails to prevent CLAD, suggesting the existence of alloimmune pathways resistant to these drugs. Here, we used mass cytometry to identify cell populations enriched in the bronchoalveolar lavage (BAL) of patients with subsequent allograft dysfunction. We show that CD4^+^CD57^+^PD1^+^ T cells emerge in stable lung transplant recipients in the first year post-transplant, conferring heightened risks for CLAD and death or re-transplantation. CD4^+^CD57^+^PD1^+^ T cells display features of senescence and secrete inflammatory cytokines. Cellular indexing by transcriptomes and epitopes (CITE-Seq) on BAL CD4^+^ T cells revealed the existence of 2 oligoclonal CD57^+^ subsets with putative cytotoxic and follicular helper functions. Finally, we observed that CD4^+^CD57^+^PD1^+^ T cells are associated with lung allograft fibrosis in a mouse model and in human explanted CLAD lungs, where they localize near airway epithelium and B cells. Together, our findings reveal the existence of an inflammatory T cell population that predicts future lung allograft dysfunction and may represent a rational therapeutic target in lung transplant recipients.

---

The study of allograft rejection has led to important advances in our understanding of cellular immunity^1^. Unfortunately, solid organ transplantation continues to be hampered by high rates of chronic rejection and graft failure despite the availability of potent immunosuppressive drugs. In lung transplantation (LT), which remains the only life- prolonging therapy for many cases of end-stage lung disease, median graft survival is only 6 years^2^ due to chronic lung allograft dysfunction (CLAD). CLAD is the result of a progressive scarring process due to accumulated injuries resulting from alloreactivity and innate immune activation caused by donor brain death, ischemia-reperfusion injury, post-transplant infections, gastroesophageal reflux, and air pollution^3^. Both acute T cell- mediated rejection and antibody-mediated rejection are well-recognized CLAD risk factors^4,5^.

LT recipients often grapple with episodes of acute lung allograft dysfunction (ALAD)^6^. We defined ALAD as a marked decrement in lung function – quantified by a decline in the forced expiratory volume in one second (FEV_1_) by 10% or more from its prior value. Despite exhaustive history, physical examination, and diagnostic testing including thoracic imaging and bronchoscopy, the cause of ALAD often is not definitively identified. Recurring ALAD events may be the result of rejection or uncontrolled alloimmunity that cannot be identified due to the poor diagnostic accuracy of transbronchial biopsies ^7,8^.

Bronchoalveolar lavage (BAL) is a technique in which saline instilled via a bronchoscope wedged in a pulmonary segmental airway is aspirated into a collection trap. BAL permits microbiologic and cytologic analysis of the distal lung compartments including the small airways and alveolar spaces. We have been studying cellular elements of the BAL using advanced multiparametric analytical approaches for the last several years^6,9–13^. Using single-cell RNA sequencing, we recently identified specific alveolar macrophage populations associated with ALAD episodes and CLAD^6^, which may represent both a mechanistic link between inflammatory insults such as infection and rejection, as well as a potential target for therapeutic intervention.

Prior studies by several groups have examined specific BAL T cell populations in LT recipients. Early work examined the question of whether changes in the CD4:CD8 ratio among BAL T cells might differentiate infection from rejection^14^. Later studies examined whether T cell activation^15^, differentiation^16^, regulatory T cells^17^ and apoptosis^18^ could distinguish CLAD, acute rejection and infection^19^. High variability between LT recipients meant that such distinctions were difficult^19^ and not useful in an individual patient. Moreover, since acute rejection and infection and CLAD can all coexist, and there are large overlaps in the distributions of pre-specified BAL immune cell populations between these clinical entities, these hypothesis-directed analyses have not led to the identification of useful biomarkers or mechanistic insights.

Here, we used mass cytometry^20^ – a high-dimensional technique in which the expression of up to 40 cell-associated proteins can be analyzed simultaneously, coupled with unbiased differential discovery approaches – to identify CD4^+^CD57^+^PD1^+^ T cells as a predictor of incident ALAD and CLAD. Apart from their usefulness as a biomarker, our data implicate CD4^+^CD57^+^PD1^+^ T cells in the mechanism of allograft loss. We find the cells in chronically rejected human and mouse lung allografts, show that they are clonally expanded, express transcripts associated with cytotoxic and helper functions, and they secrete inflammatory cytokines. Further, they are senescent, hypoproliferative, and resistant to cell death *in vitro.* Our observations therefore suggest the existence of a specific alloimmune pathway that can escape conventional immunosuppression that may represent a valuable therapeutic target.

## Results

### Identification of a T cell subset associated with lung allograft dysfunction

Early detection of alloimmune responses in the graft – before ALAD onset – is likely to provide both important prognostic information as well as an opportunity for early intervention. In pursuit of a greater understanding of ALAD mechanisms, we prospectively collected and viably cryopreserved BAL cells from 50 Toronto Lung Transplant Program (TLTP) LT recipients presenting for surveillance bronchoscopy at 3 months post-transplant, and obtained two more follow up samples from each recipient to form a longitudinal cohort (**Figure 1A; Supp Fig 1**). We performed mass cytometry^20^ using a panel of 36 heavy metal-conjugated antibodies **(Supp Table 1)**. Conventional flow cytometric gating revealed considerable inter- and intra-patient heterogeneity in BAL cell composition (data not shown). Unsupervised multidimensional clustering using FlowSOM ^21^ demonstrated similar variation (**Figure 1B, Supp Fig 2**). In order to facilitate identification of cell populations associated with allograft dysfunction, we randomly divided the cohort into discovery and validation subsets (n=25 each; **Supp Table 2**). Longitudinal samples in the discovery group were then used to search for differences in the BAL cellular composition between stable patients and those experiencing ALAD within 30 days of each bronchoscopy. Initially we employed cluster identification, characterization and regression (CITRUS^22^) on all BAL T cells from all the time-points for all patients in the cohort. This analysis revealed that CD4^+^CD57^+^PD1^+^ T cells were the top differentially represented cell population between patients with and without ALAD within 30 days of at least one bronchoscopy **(Supp Fig 3**). CITRUS does not adequately account for the paired design of our cohort (multiple samples per patient), nor does it incorporate time as a factor in the analysis. We therefore also applied a generalized linear mixed model (GLMM) pipeline described for CyTOF analysis^23^. To facilitate this analysis, we used FlowSOM to identify 20 clusters of T cells (**Figure 1C**). T cell subsets with greater representation in samples associated with ALAD compared to samples associated with stability were identified using this approach (**Figure 1D**). The expression profile of T cell markers revealed that the CD4^+^ CD57^+^PD1^+^ T cell population was significantly differentially represented in ALAD (**Figure 1E, Supp Fig 4**). Collectively, these data suggested that there is a strong association between CD4^+^CD57^+^PD1^+^ T cells and incident ALAD.

**Figure 1.**
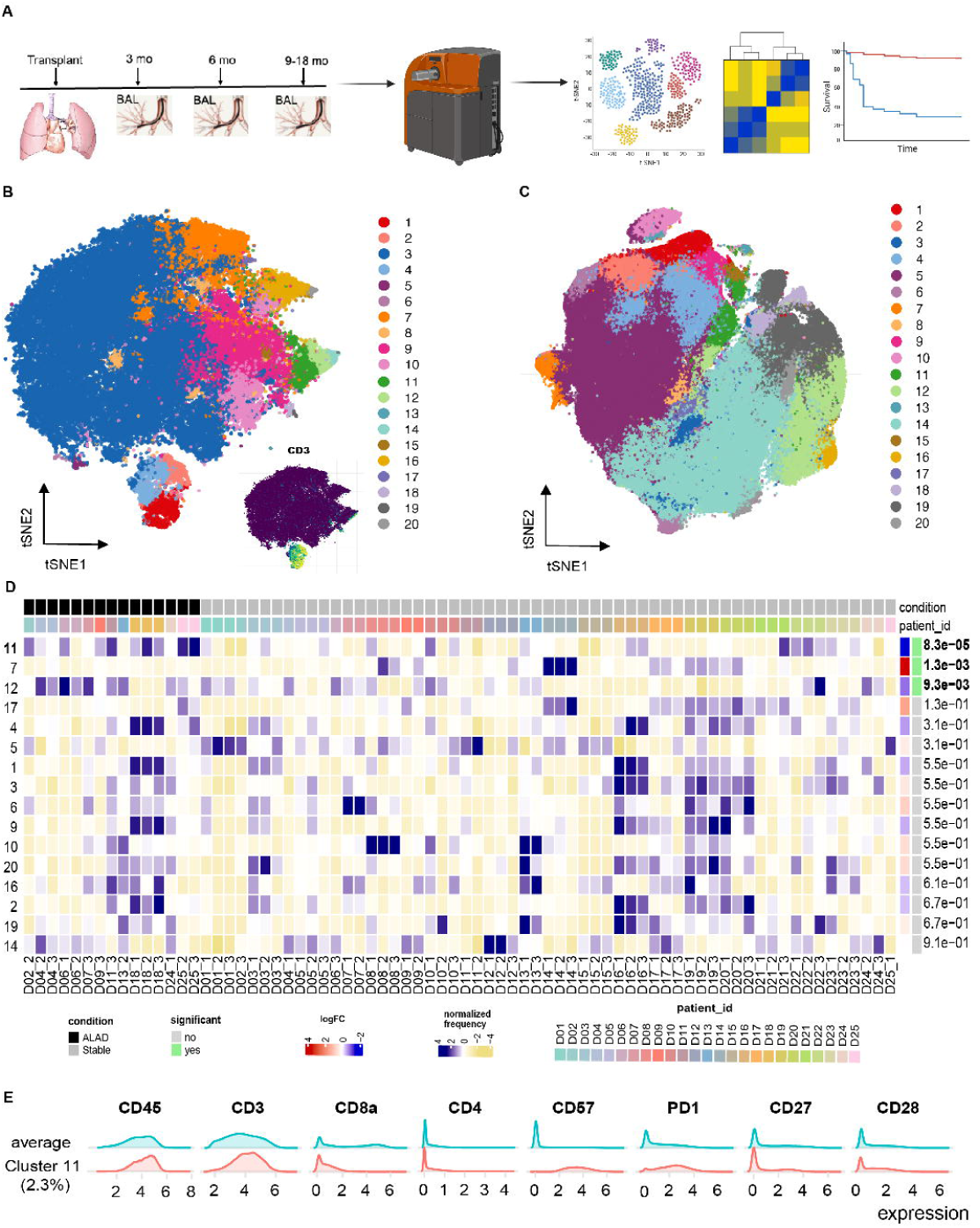
Identification of a BAL CD3^+^ T cell population associated with lung allograft dysfunction. **A.** Study schematic. Fifty consecutive patients presenting for bronchoscopy at 3 months post-transplant were enrolled. For each enrolled patient, BAL cells at three time points (3mo, 6mo, 9mo or next available) were collected. BAL cells were cryopreserved and subjected to mass cytometry (see supplementary table 1); all samples from each patient were analyzed in the same mass cytometry experiment. **B.** t- distributed stochastic neighbour embedding (tSNE) plots of BAL CD45^+^ cells falling into 20 clusters (highlighted in different colours) identified by FlowSOM. Inset tSNE plot highlighting expression of CD3^+^ T cells is shown at lower right. **C.** t-SNE plot based on arcsinh-transformed marker expression in 500 randomly selected cells per sample from the discovery set (n=25, all 3 time points shown). Cells are coloured according to the 20 cell populations obtained with FlowSOM after the metaclustering step with ConsensusClusterPlus. **D.** Differential analysis heatmap of arcsine-square-root transformed cell frequencies that were subsequently normalized per cluster (rows, 20 clusters identified in the panel C). Statistically significant clusters are highlighted by green boxes at right along with p-values. **E.** Analysis of molecules expressed by all cells (average) compared with those expressed by the most differentially represented cell population in ALAD patients (cluster 11) reveals that the latter is CD45^+^CD3^+^CD4^+^CD57^+^PD1^+^.

**Figure 2.**
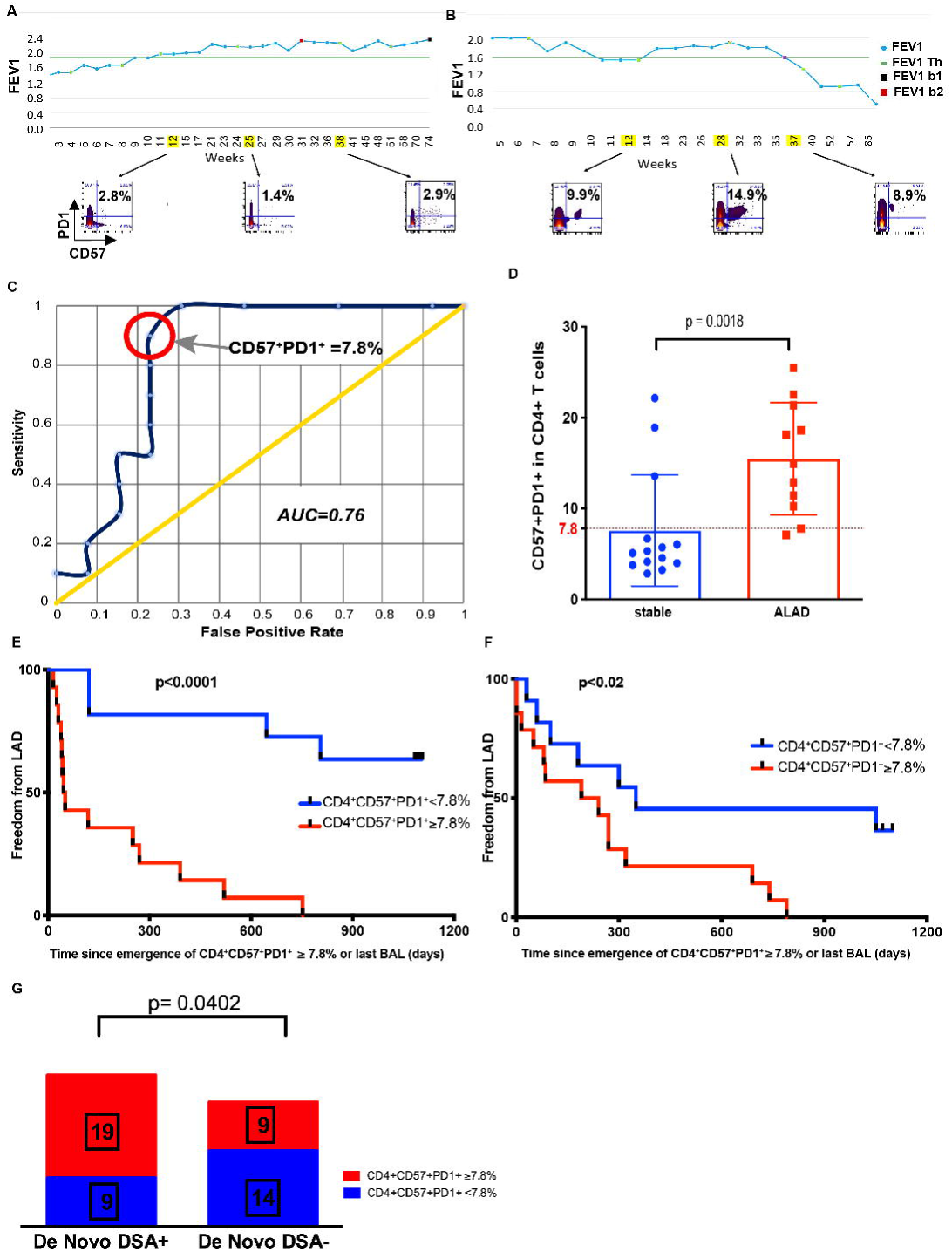
Relationship of BAL CD4^+^CD57^+^PD1^+^ T cells with incident allograft dysfunction. Representative examples of forced expiratory volume in one second (FEV1) over time in weeks post-transplant for stable (**A**) and ALAD (**B**) lung transplant recipients in the discovery cohort. Weeks during which bronchoscopy was performed are highlighted in yellow; scatterplots showing expression of CD57 and PD1 in the CD4^+^ T cell compartment of the BAL are shown for each time point. Green line indicates 80% of best post-transplant baseline, below which CLAD may be diagnosed. **C.** Receiver- operator characteristic (ROC) curve analysis of the performance characteristics of CD57^+^PD1^+^ cells as a proportion of the BAL CD4^+^ compartment for diagnoses of subsequent ALAD. The area under the curve (AUC) = 0.76. The percentage of CD57^+^PD1^+^ T cells providing optimal discrimination of ALAD and stable patients (7.8%) is circled in red. **D.** The highest proportion of CD57^+^PD1^+^ T cells observed in all BAL samples from each patient (discovery cohort, n=25) is shown, divided by the presence or absence of ALAD within 30 days. Mann-Whitney test p=0.0018. Dotted line shows the 7.8% threshold identified in ROC curve in C. **E.** Time to lung allograft dysfunction from the emergence of CD57^+^PD1^+^ cells ≥ 7.8% of CD4^+^ T cells in the BAL (red line), or from the final BAL analyzed (blue line) in the discovery cohort (n=25). Log rank test p < 0.0001. **F.** Time to lung allograft dysfunction from the emergence of CD57^+^PD1^+^ cells ≥ 7.8% of CD4^+^ T cells in the BAL (red line), or from the final BAL analyzed (blue line) in the validation cohort (n=25). Log rank test p < 0.02. G. Of 28 patients who developed de novo DSA, 19 (67.9%) had CD57^+^PD1^+^ cells ≥ 7.8% of CD4^+^ T cells in at least one BAL sample; only 9 of 23 (39.1%) patients who did not develop de novo DSA had CD57^+^PD1^+^ cells ≥ 7.8% of CD4^+^ T cells in at least on BAL sample. Chi square test p = 0.0402.

**Figure 3.**
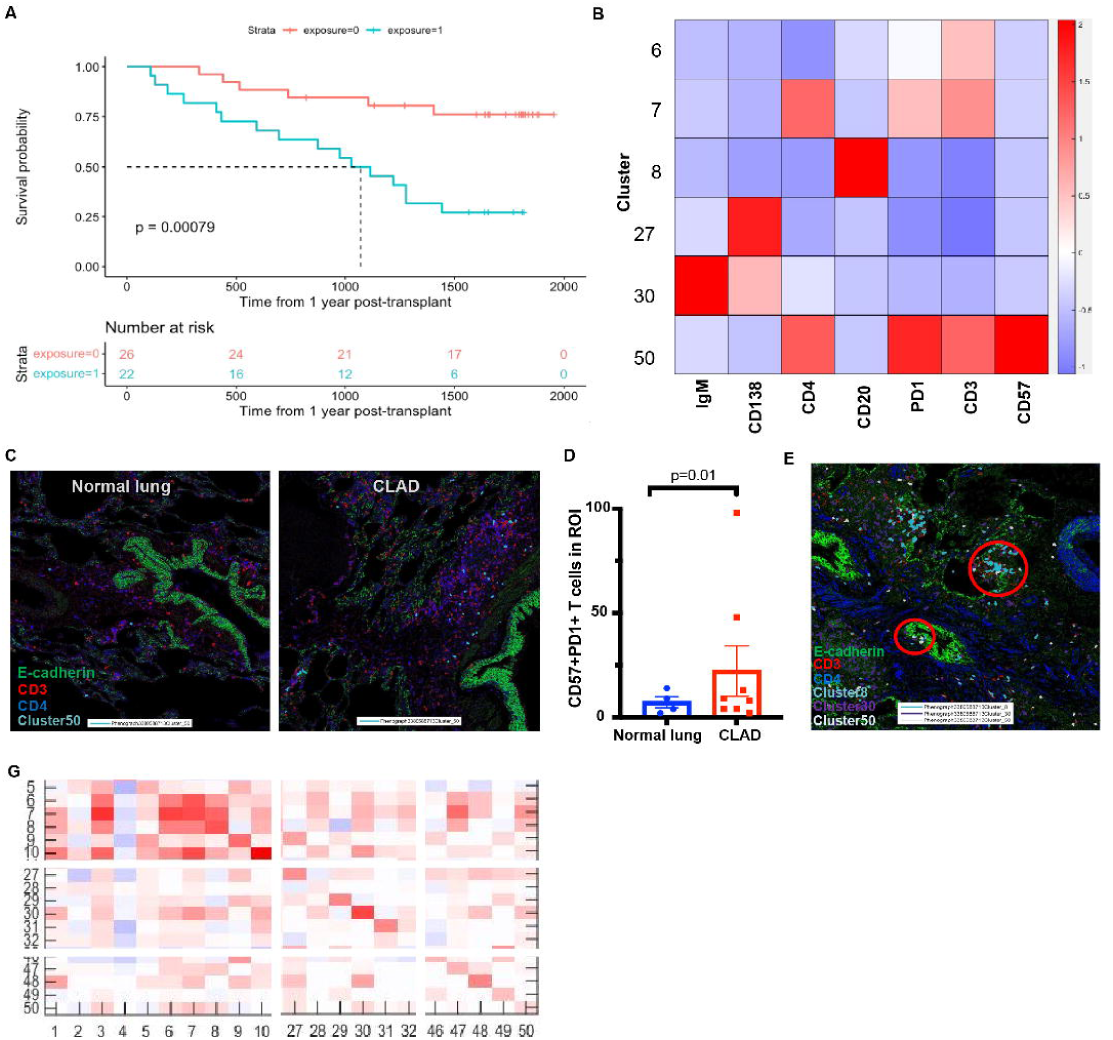
CD4^+^CD57^+^PD1^+^ T cells are associated with an increased risk of death or re-transplantation and are found in CLAD lung tissue. **A.** Kaplan-Meier curves showing time to death or re-transplantation from 12 months post-transplant. Log-rank test p = 0.00079. **B.** Selected heatmap of protein expression for selected clusters and markers to highlight protein profile of cluster 50. **C.** Imaging mass cytometry of normal donor lung tissue (left) and CLAD lung tissue obtained at the time of re-transplantation (right). E-cadherin^+^ epithelial cells (green pseudocolour), red CD3^+^ T cells (red pseudocolour), CD4^+^ cells (blue pseudocolour), and CD4^+^CD57^+^PD1^+^ T cells (cluster 50, pale blue pseudocolour) are shown. **D.** CD4^+^CD57^+^PD1^+^ T cells are more numerous in CLAD lung tissue. Data from 3 regions of interest (ROIs) from each of n=8 CLAD, and n=4 normal lungs are shown. **E.** In CLAD lung tissue, CD4^+^CD57^+^PD1^+^ T cells (cluster 50, white) are found in association with airways (green pseudocolour E-cadherin^+^ cells) and with two distinct B cell populations (cluster 8, pale blue pseudocolour and cluster 30, purple pseudocolour). **F.** Cell-cell interaction heatmap shows association of CD4^+^CD57^+^PD1^+^ T cells (cluster 50) with epithelial cells (cluster 27) and B cells (clusters 8 and 30). G. Liquid phase mass cytometry performed on single cell suspensions obtained from CLAD lung tissue show variable presence of CD4^+^CD57^+^PD1^+^ T cells.

**Figure 4.**
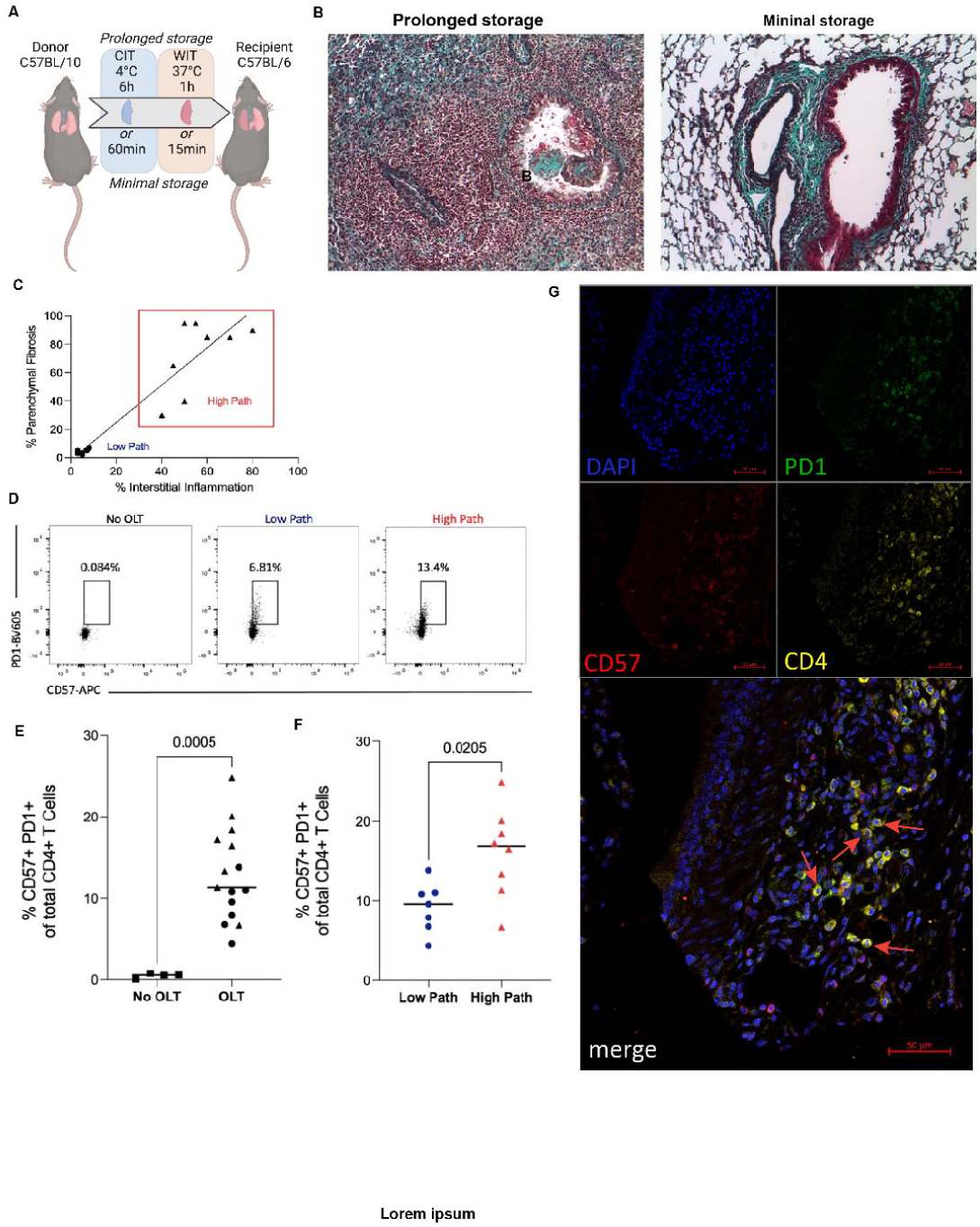
CD4^+^CD57^+^PD1^+^ T cells are associated with fibrosis in a mouse model of chronic lung allograft rejection. **A.** C57BL/10 (B10) donor lungs were subjected to either prolonged (6h cold ischemic time (CIT) at 4°C followed by 1h warm ischemic time (WIT) at 37°C) or minimal (60min CIT followed by 15min WIT) storage prior to orthotopic left lung transplantation into minor alloantigen-mismatched C57BL/6 (B6) recipients. **B.** After 28 days, paraffin-embedded sections of lung tissue were subjected to Masson trichrome staining (200x magnification). **C.** Grafts were scored using a published scoring system^28^ that quantifies the percentage of the graft affected by interstitial inflammation and parenchymal fibrosis. Grafts with high-grade pathology (High Path) are represented as triangles, and those with minimal pathology (Low Path) are represented as circles. **D.** Single-cell suspensions of naïve B6 lungs (No OLT) and lung allografts obtained at 28 days post-transplantation were subjected to flow cytometry. Analyses are gated on CD4^+^ T cells. Percentages of cells in the CD57^+^PD-1^+^ gate are shown in the representative plots. **E.** Percentage of CD4^+^ T cells expressing CD57 and PD-1 is shown for naïve B6 lungs (No OLT, squares), Low Path grafts (circles) and High Path grafts (triangles). Student’s t test, p=0.0005. **F.** Percentage of CD4^+^ T cells expressing CD57 and PD-1 is shown for Low Path grafts (circles) and High Path grafts (triangles). Student’s t test, p=0.0205. **G.** Immunofluorescence of prolonged storage, High Path lung allograft showing PD1 (green), CD57 (red), CD4 (yellow) and DAPI (blue) and merged image.

### CD4^+^CD57^+^PD1^+^ T cells appear in the BAL prior to the onset of allograft dysfunction

Next, we assessed the relationship of BAL CD4^+^CD57^+^PD1^+^ T cells to ALAD onset. We observed that while patients with stable lung allograft function had low frequencies of CD57^+^PD1^+^ cells in the BAL CD4^+^ T cell compartment (**Figure 2A**), these cells typically accumulated in higher frequencies prior to the onset of ALAD (**Figure 2B**). Receiver-operating characteristic (ROC) curve analysis showed that having ≥ 7.8% CD57^+^PD1^+^ cells in the BAL CD4^+^ T cell compartment optimally discriminated concurrent ALAD from stable patients, with an area under the ROC curve of 0.76 (**Figure 2C**). Using this criterion, 10 of 11 (91%) LT recipients with ALAD within 30 days of any of their bronchoscopies demonstrated ≥ 7.8% CD57^+^PD1^+^ in the CD4^+^ T cell compartment of at least one of their BAL samples; in contrast, only 3 of 14 (21%) stable LT recipients had CD57^+^PD1^+^ cells exceeding this threshold (**Figure 2D**). Moreover, having this biomarker conferred a reduced time-to-ALAD in univariable survival analysis (**Figure 2E**). We then applied the 7.8% threshold to the validation subset (n=25) and observed a similar reduction in time-to-ALAD from first appearance of the cells in LT recipients with at least one BAL sample in which it was exceeded, compared to LT recipients who did not have any BAL samples in which the threshold was exceeded (**Figure 2F**). Together, these data illustrated that in LT recipients developing subsequent allograft dysfunction, CD57^+^PD1^+^ cells frequently exceeded the 7.8% threshold in BAL samples obtained prior to ALAD and CLAD **(Supp Fig 5).**

**Figure 5.**
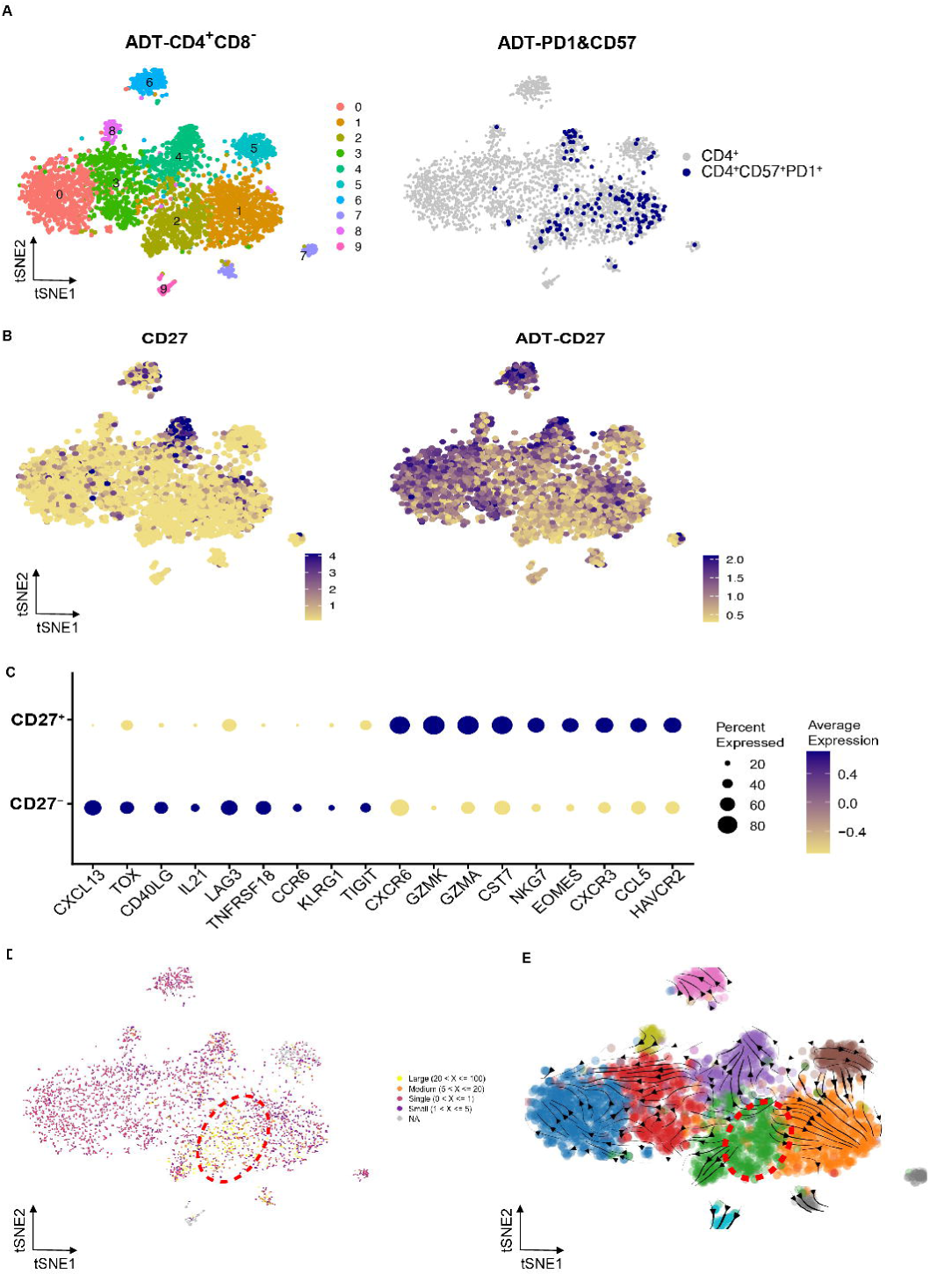
Identification of distinct CD27^+^ and CD27^-^ subsets of CD4^+^CD57^+^PD1^+^ T cells. CITE-Seq was performed on 4 BAL samples from patients with ALAD. **A.** After downsampling and integration, BAL CD4^+^ T cells (CD3^+^CD4^+^CD8^-^ identified by antibody-derived oligonucleotide tag, ADT) were displayed on a tSNE plot revealing 10 distinct cell clusters (left tSNE plot). Right panel displays CD57 and PD-1 expression (blue dots) on a feature plot showing all BAL CD4^+^ cells. **B.** tSNE plot demonstrating CD27 RNA expression (left) and at the protein level (ADT-CD27, right). **C.** Dot plot of the selected top differentially expressed genes in BAL CD27^+^ and CD27^-^ CD4^+^CD57^+^PD-1^+^ T cells. Purple colour indicates higher expression; dot size reflects the percentage of cells in each population expressing the gene. **D.** tSNE plot overlaid with colours reflecting TCR clonal expansion. Dotted red circle indicates highly expanded clones within the CD27^-^ population.. **E.** tSNE plot overlaid with RNA velocity trajectories (arrows). Dotted red circle shown in D is overlaid here indicating the likely origin of CD4^+^CD57^+^PD-1^+^ T cells.

Anti-human leukocyte antigen donor-specific antibodies (DSA) are an adverse prognostic factor in lung transplantation^24^. Patients with ≥7.8% CD57^+^PD1^+^ cells in the BAL CD4^+^ T cell compartment were more likely to develop de novo DSA than those without (**Figure 2G**). BAL CD4^+^CD57^+^PD1^+^ T cells were not simply a surrogate for other conventional clinical and bronchoscopic parameters, since clinical symptoms, biopsy-proven vascular or airway rejection, BAL pathogens and antimicrobial treatments did not differ significantly between patients with and without ≥ 7.8% CD57^+^PD1^+^ cells in the BAL CD4^+^ T cell compartment **(Supp Table 3)**. Similarly, although the number of events was small, having CD57^+^PD1^+^ cells above the 7.8% threshold was not clearly associated with an immediate post-bronchoscopy augmentation of immunosuppression or change in calcineurin inhibitor **(Supp Table 4)**. Taken together, these observations indicate that BAL CD4^+^CD57^+^PD1^+^ T cells may be a useful biomarker of incident lung allograft dysfunction that adds significant diagnostic utility to conventional clinical monitoring.

### CD4^+^CD57^+^PD1^+^ T cells confer an increased risk for CLAD and death and are present in chronically rejected human lung allografts

To examine whether the presence of CD4^+^CD57^+^PD1^+^ T cells in the BAL samples of LT recipients increase the risk for CLAD, we followed all 50 individuals in our cohort for up to 2000 days post-LT (median follow-up 1788 days). Using the appearance of ≥7.8% CD57^+^PD1^+^ cells in the BAL CD4^+^ T cell compartment or the last analyzed BAL as the index event, we found that time to CLAD was significantly shorter in those patients whose BAL samples exceeded the 7.8% threshold (log rank p=0.0071, **Supp Figure 6**). We then performed Cox proportional hazards analysis, excluding two patients who developed CLAD prior to their final BAL samples (**Supp Fig 1**). Treating the appearance of ≥7.8% CD57^+^PD1^+^ cells in the BAL CD4^+^ T cell compartment as a time-dependent covariate, Cox proportional hazards analysis revealed a hazard ratio (HR) of 3.61 (95% CI 1.55-8.39) for time-to-CLAD. In this and other survival analyses, patients were followed until 31 December 2022 (median follow up time from transplantation: 2245 days). The increased risk for CLAD associated with CD4^+^CD57^+^PD1^+^ cells persisted in bivariable models adjusted for age at transplant (HR 3.67, 95% CI 1.57-8.55), sex (HR 3.45, 95% CI 1.47-8.07), and primary cytomegalovirus (CMV) mismatch (donor seropositive, recipient seronegative; HR 4.02, 95% CI 1.70-9.52).

**Figure 6.**
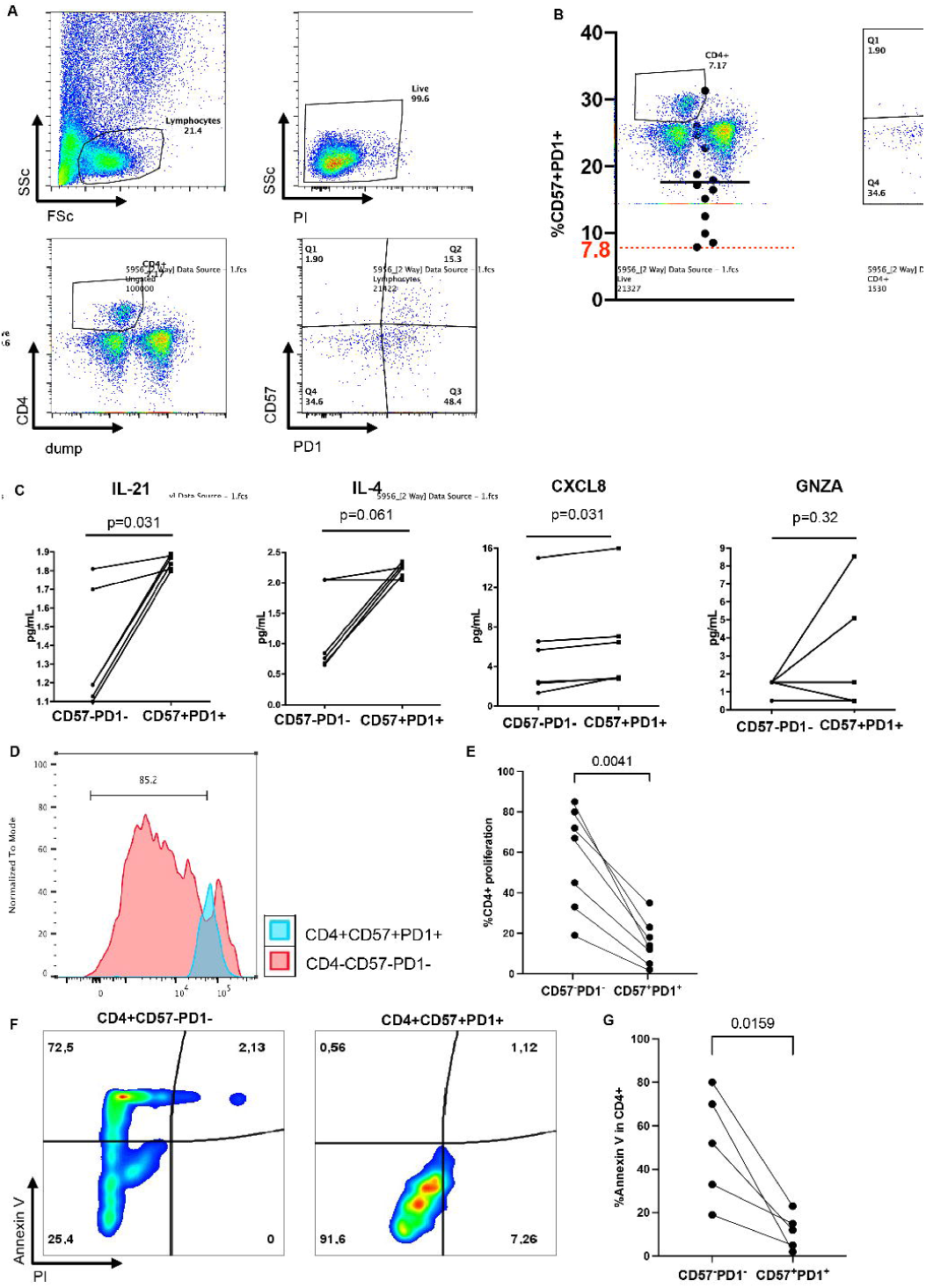
Functional properties of CD4^+^CD57^+^PD1^+^ T cells. **A.** CD4^+^CD57^+^PD1^+^ T cells can be identified by conventional flow cytometry. BAL cells were stained with CD4, CD57 and PD1 antibodies and CD57^+^PD1^+^ cells were identified within the CD4^+^ population, allowing quantification and cell sorting. **B**. Percentage of CD57^+^PD1^+^ cells within the CD4^+^ gate is shown for n=14 patients with ALAD. **C.** Sorted CD57^+^PD1^+^ and CD57^-^PD1^-^ CD4^+^ T cells from ALAD BAL samples were stimulated *in vitro* with anti- CD3/anti-CD28 antibodies overnight. Culture supernatants were subjected to multiplexed cytokine quantification. IL-21 (p=0.031), IL-4 (p=0.062), CXCL8 (IL-8, p=0.031) and GZMA (p=0.062) were generally secreted in greater concentrations by CD57^+^PD1^+^ T cells than their CD57^-^PD1^-^ counterparts, Wilcoxon matched-pairs signed rank tests. **D.** Sorted CD57^+^PD1^+^ (blue) and CD57^-^PD1^-^ CD4^+^ (pink) T cells from ALAD BAL were examined for proliferation in response to CD3/CD28 stimulation by dilution of CFSE. **E.** The percentage of CD57^+^PD1^+^ T cells undergoing proliferation was lower compared to that of their CD57^-^PD1^-^ counterparts. Linked CD57^+^PD1^+^ and CD57^-^PD1^-^ cells are sorted from same LT recipient. Wilcoxon matched pairs signed rank test, p=0.0041. **F.** Sorted CD57^+^PD1^+^ T cells (right) and their CD57^-^PD1^-^ counterparts (left) were subjected to RICD. Annexin V and propidium iodide (PI) staining is shown. **G.** Annexin V staining was lower in CD57^+^PD1^+^ T cells compared to their CD57^-^PD1^-^ counterparts. Linked CD57^+^PD1^+^ and CD57^-^PD1^-^ T cells are sorted from same LT recipient. Wilcoxon matched pairs signed rank test, p=0.0159.

Similarly, a Cox proportional hazards model revealed that ≥7.8% CD57^+^PD1^+^ cells in the BAL CD4^+^ T cell compartment was associated with a HR of 4.82 (95% CI 1.8-13.1) for time-to-death or re-transplantation. The effect persisted in bivariable models adjusting for age at transplant (HR 4.82, 85% CI 1.8-13.1), sex (HR 4.84, 1.8-13.2) and primary CMV mismatch (HR 4.92, 95% CI 1.8-13.4). These time-to-CLAD and time-to-death or re-transplantation analyses are presented in **Supp Table 5**. We then examined time-to-death or re-transplantation in a survival analysis landmarked at 12 months post-transplant. LT recipients with ≥ 7.8% CD57^+^PD1^+^ cells in the BAL CD4^+^ T cell compartment had an increased risk of death or re-transplantation (p=0.00079, **Figure 3A**).

These observations suggested that CD4^+^CD57^+^PD1^+^ T cells may lie on the mechanistic pathway leading to lung allograft fibrosis. Therefore, we sought to identify them in CLAD lung tissue obtained at the time of re-transplantation^25^ using imaging mass cytometry (IMC^26^). CLAD and normal lung tissue sections were stained with a panel of 35 heavy metal conjugated antibodies **(Supp Table 6)** and subjected to IMC. Unsupervised clustering identified CD4^+^CD57^+^PD1^+^ T cells as one of the cell populations present (cluster 50 in **Figure 3B**), with more cells in CLAD compared to normal tissue (**Figure 3C-D**). Image reconstruction and proximity analysis revealed that CD4^+^CD57^+^PD1^+^ T cells were frequently located adjacent to distinct B cell populations, including trafficking B cells (CD20^+^CCR6^+^, Cluster 8) and plasma cells (IgM^+^CD138^+^, cluster 30) and airways (**Figure 3E-F**). The enrichment of CD4^+^CD57^+^PD-1^+^ T cells in CLAD tissue compared to normal lung tissue may indicate their involvement in disease development.

### CD4^+^CD57^+^PD1^+^ T cells infiltrate mouse lung allografts and associate with the severity of chronic rejection

We have developed a minor alloantigen-mismatched mouse orthotopic lung transplant model in which a minimal graft storage period prior to transplantation results in mild or absent allograft fibrosis at 28 days post-transplant, while prolonged warm and cold ischemic storage prior to transplantation leads to B cell- and alloimmune-dependent fibrosis with features of human chronic lung allograft rejection^27^ (**Figure 4A-B**). We asked whether CD4^+^CD57^+^PD-1^+^ T cells might be found within lung allografts in this model, and whether their abundance correlated with the degree of allograft fibrosis. Allografts were scored in blinded fashion using our published scoring system^28^, which includes assessment of the proportion of the allograft affected by interstitial inflammation and fibrosis. As we have observed previously^27,28^, pathology was variable, with some animals exhibiting higher degrees of fibrosis and inflammation (**Fig 4C**). Flow cytometry performed on single-cell suspensions of digested lung allograft tissue revealed that CD57^+^PD1^+^ T cells were common among lung allograft CD4^+^ T cells, but rare in normal lung tissue (**Figure 4D**). It also showed that allografts with higher degrees of pathology – as determined by the presence of both interstitial inflammation and parenchymal fibrosis – had higher frequencies of CD57^+^PD1^+^ T cells compared to those with a low degree of pathology (**Figure 4E-F**). Further, CD4^+^CD57^+^PD1^+^ T cells were readily detectable in allograft sections by immunofluorescence (**Figure 4G**). Together, these data support the concept that CD4^+^CD57^+^PD1^+^ T cells accumulate in rejecting lung allografts and correlate with the severity of inflammation and fibrosis in mice.

### Identification of two transcriptionally distinct CD4^+^CD57^+^PD1^+^ cell subsets using CITE-Seq

To characterize gene expression in CD4^+^CD57^+^PD1^+^ T cells, we enriched CD3^+^ T cells from four ALAD BAL samples and performed cellular indexing by transcriptomes and epitopes (CITE-Seq^29^) with 5’ end sequencing. We applied a panel of oligonucleotide-conjugated antibodies, including CD4, CD57 and PD1, to ALAD BAL samples containing at least 7.8% CD57^+^PD1^+^ cells in the CD4^+^ T cell compartment as determined by flow cytometry. The conjugated oligonucleotides or antibody-derived tags (ADTs) identified in CITE-Seq allowed confident linkage of transcriptomes with protein expression.

BAL CD4^+^ T cells from patients with ALAD fell into 9 subclusters in multidimensional space (**Figure 5A**). Based on CD57 and PD1 protein expression, CD4^+^CD57^+^PD1^+^ T cells assembled into two main areas, cluster 4 and the closely adjacent clusters 1 and 2 **(Figure 5A, Supp Fig 7A-B**). We noted that CD27 gene and protein expression was enriched in cluster 4 compared to clusters 1 and 2 (**Figure 5B).** The CD27^-^ population had a gene expression profile similar to peripheral helper T (Tph) cells described in rheumatoid arthritis^30^; differentially expressed genes in this population included CXCL13, TOX, IL-21, CD40L, TIGIT, ICOS and TNFSRF18 **(Figure 5C, Supp Fig 7C**). These cells are characterized by the absence of the conventional follicular helper T cell markers CXCR5 and Bcl6, and by expression of the chemokine CXCL13 ^30,31^.

Accumulating studies have shown that Tph cells are involved in various acute and chronic inflammatory diseases ^30,32^, and can promote B cell differentiation into plasmablasts through interactions involving IL-21 and SLAMF5 ^30^. Therefore, the Tph-like features of CD27^-^ CD4^+^CD57^+^PD1^+^ T cells, along with their proximity to B cells (**Figure 3E-F**), their association with de novo DSA (**Figure 2G**) suggest that they may provide B cell help.

The CD27^+^ population expresses several cytotoxicity-related transcripts including granzymes and NKG7^33^ (**Figure 5C**). They also express CXCR6, which has been associated with tissue resident memory CD8^+^ T cell localization to airways^34^ and is in keeping with our observation of CD4^+^CD57^+^PD1^+^ T cells in proximity to airways (**Figure 3F**).

We next examined TCR CDR3 sequences in the CITE-Seq data, which revealed that CD57^+^PD1^+^CD27^-^ cells are clonally expanded to a greater extent than CD57^+^PD1^+^CD27^+^ and other BAL CD4^+^ T cells (**Figure 5D**). This suggested that they had recently undergone antigen-driven expansion in the lung. Analysis of the top 20 expanded TCR clonotypes did not infer any known antigen. However, the high similarity among the top 20 expanded TCRs may suggest that they may be alloreactive **(Supp Figure 8**). Pseudotime trajectory analysis revealed that CD57^+^PD1^+^CD27^-^ cells and their CD27^+^ counterparts may share a common origin (**Figure 5E**). Together, the TCR sequence data and pseudotime analysis suggested that BAL CD4^+^CD57^+^PD1^+^ T cells either differentiate into Tph or cytotoxic effector cells. The former have undergone significant clonal expansion, whereas the latter have a distinct TCR repertoire and less clonal expansion, suggesting that they may be responding to different antigens in the lung microenvironment.

Finally, both CD27^-^ and CD27^+^ subsets demonstrated expression of previously defined senescence and exhaustion transcripts including TIGIT, KLRG1, LAG3, HAVCR2/TIM-3 and EOMES which have been described in different acute and chronic inflammatory conditions^35–37^ . This observation is in keeping with reports that have described different senescent CD57^+^ and/or PD1^+^ T cell populations in other disease states ^38,39^.

### Functional analysis of CD4^+^CD57^+^PD1^+^ T cells reveals features of inflammation and senescence

Considering the potential distinct functional properties – Tph, cytotoxic, and senescent – of CD4^+^CD57^+^PD1^+^ T cells, we set out to characterize their functions *in vitro*. For this purpose, CD4^+^ T cells from BAL samples of 15 lung transplant recipients experiencing ALAD were sorted to obtain CD4^+^CD57^+^PD1^+^ T cells and their CD4^+^CD57^-^PD1^-^ counterparts (**Figure 6A**). Fluorescence-activated cell sorting data demonstrated that in all cases, CD57^+^PD1^+^ cell frequency was ≥7.8% of BAL CD4^+^ T cells (**Figure 6B**). Based on the hypothesis that CD4^+^CD57^+^PD-1^+^ T cells may be senescent, we predicted that sorted cells would exhibit decreased proliferation, enhanced cytokine secretion, and resistance to TCR restimulation-induced cell death (RICD) when compared to CD57^-^ PD1^-^ cells.

Supernatants from cells stimulated overnight were collected and subjected to a multiplexed cytokine assay. This revealed that CD4^+^CD57^+^PD1^+^ cells – compared to CD4^+^CD57^-^PD1^-^ cells sorted from the same BAL samples – expressed more IL-21, IL-4, CXCL8 and granzyme A (**Figure 6C**). In keeping with our CITE-Seq data, these findings suggested that CD4^+^CD57^+^PD1^+^ T cells may have Tph or cytotoxic properties – or both. Stimulated CD4^+^CD57^+^PD1^+^ T cells and their CD57^-^PD1^-^ counterparts were then tested for proliferation, based on dilution of CFSE (**Figure 6D).** Proliferation data revealed that CD4^+^CD57^+^PD1^+^ T cells exhibited significantly less proliferation compared to CD4^+^CD57^-^PD1^-^ cells (**Figure 6E**), in keeping with T cell senescence ^38,39^. Finally, CD4^+^CD57^+^PD1^+^ T cells were less susceptible to RICD compared to CD4^+^CD57^-^PD1^-^ cells (**Figure 6F, G)**. Together these data imply that CD4^+^CD57^+^PD1^+^ T cells are senescent but pro-inflammatory T cells that are resistant to antigen-induced death. These observations may help to explain the emergence of ALAD and CLAD in lung transplant recipients despite potent triple-drug immunosuppression, which would be predicted to be less effective against these cells.

## Discussion

In this study, we used a high dimensional mass cytometric discovery approach to identify a T cell population in lung transplant recipient BAL that is associated with a heightened risk for future allograft dysfunction, failure, and death. The risk conferred by the appearance of CD4^+^CD57^+^PD1^+^ T cells in the BAL remained elevated after bivariate adjustments for age, sex, and donor and recipient CMV serostatus mismatch.

In keeping with previous work examining lung transplant recipient BAL, our samples contained a diversity of leukocyte populations. The prior studies demonstrated statistically significant relationships between BAL CD8^+^ T cells, NK cells, eosinophils, Tregs and outcomes in lung transplantation, but this has not led to meaningful improvements in clinical care because wide overlaps in the frequency distributions of these populations have meant that they lack discriminatory power in individual patients^14,15,19,40–43^. In contrast, while CD4^+^CD57^+^PD1^+^ T cells will require further validation in larger studies, their prevalence among CD4^+^ T cells discriminated patients who remained stable from those about to undergo ALAD with high precision, in a discovery cohort and confirmed in an independent validation cohort. Our observation therefore represents a promising biomarker that may, in the future, permit the selection of lung transplant recipients for targeted preventative interventions.

A review of clinical and laboratory variables present at the time of each bronchoscopy did not reveal any clear association between the appearance of ≥ 7.8% CD57^+^PD1^+^ T cells in the BAL CD4^+^ T cell compartment and other indicators of lung allograft inflammation or dysfunction. This observation likely reflects the relatively poor performance of histopathologic and cytologic findings in the diagnosis of lung allograft rejection. Indeed, interpretation of transbronchial biopsy histology is fraught, with high rates of ungradable biopsies and substantial inter-observer disagreement among pathologists^7,8^. It is therefore perhaps not surprising that bronchoscopies with high frequencies of CD4^+^CD57^+^PD1^+^ T cells showed no particular association with histologic rejection. Likewise, we found that this T cell population is not a marker of lung allograft infection, since there were no differences in pathogen isolation from the same BAL samples. Our interpretation of these findings is that analysis of BAL CD4^+^CD57^+^PD1^+^ T cells may dramatically improve the utility of surveillance bronchoscopy after LT, which will require further evaluation in larger prospective studies.

Beyond their value as a biomarker of future allograft dysfunction, several lines of evidence support the notion that CD4^+^CD57^+^PD1^+^ T cells are mechanistically relevant to the pathogenesis of chronic lung allograft rejection and fibrosis. First, our CITE-Seq data indicate that the majority of clonally expanded CD4^+^ T cell populations in the BAL of patients with ALAD were CD57^+^PD1^+^, which indicates that these cells have undergone antigen-driven expansion in the lung. Further, in addition to their appearance in BAL, we detected CD4^+^CD57^+^PD1^+^ T cells in human explanted CLAD lung tissue obtained at the time of re-transplantation. The cells were observed in proximity to B cells and small airways; coupled with the finding that the transcriptome and secretome of BAL CD4^+^CD57^+^PD1^+^ T cells includes IL-21 and granzymes, this observation suggests that they are likely participating in allograft injury and may subserve Tph functions. In support of this assertion, in our ischemia-reperfusion injury-augmented mouse model of human CLAD, we observed CD4^+^CD57^+^PD1^+^ T cells in the lung allografts of mice with marked fibrotic pathology. In contrast, allografts with milder fibrosis did not contain many CD4^+^CD57^+^PD1^+^ T cells. Interestingly, we have previously shown that fibrosis in this model is B cell dependent^27^, and so in future work it will be important to consider the possible existence of a CD4^+^CD57^+^PD1^+^ T cell-B cell axis that drives chronic lung allograft rejection in mice and humans.

Accumulating evidence has revealed an association of CD4^+^CD57^+^ T cells – with or without associated PD-1 expression – with autoimmune diseases, infectious diseases, and cancers ^44^. CD4^+^CD57^+^ T cells have been detected in the peripheral blood of kidney transplant recipients undergoing rejection ^45^, particularly in costimulation blockade resistant rejection ^46,47^. Of note, the CD4^+^CD57^+^ T cells associated with rejection in kidney transplantation were measured in the PBMC and exhibit low expression of PD1. In another study, Brugière and colleagues observed that the risk of CLAD was increased in patients with a high proportion of circulating CD4^+^CD57^+^ILT2^+^ T cells, which could be inhibited by HLA-G^48^. However, in that study, PD-1 expression was not assessed, and the association with CLAD was not as strong as we report here. Therefore, it is unclear whether any of these peripheral CD4^+^CD57^+^ populations bear a functional resemblance or developmental relationship to those that we have described within lung allografts. Moreover, although peripheral blood CD57^+^ T cells are also reported to reflect CMV infection and/or age^49^, our patients’ characteristics did not suggest any association between age and/or CMV seropositivity with CD4^+^CD57^+^PD1^+^ T cells.

PD1 expression by CD8^+^ T cells in tumor microenvironments is associated with cancer progression, and generally reflects a state of exhaustion in which effector functions are deactivated. BAL CD4^+^CD57^+^PD1^+^ T cells also expressed several additional molecules that have been associated with exhaustion, including TIM-3, LAG3 and CTLA-4. While analysis of the cells *ex vivo* revealed that they were both hypoproliferative and resistant to T cell RICD, their production of inflammatory cytokines – which is not a feature of exhaustion – suggests that they are senescent rather than exhausted. Of note, senescent CD4^+^ and CD8^+^ T cells in humans were described as having the phenotype CD45RA^+^KLRG1^+^CD57^+^CD28^-^CD27^-^^50,51^. Animal and human data indicate that senescent T cells participate in other chronic inflammatory, degenerative and metabolic disorders^39^.

Lung transplant recipients typically receive immunosuppressive drugs – calcineurin inhibitors, corticosteroids, and antimetabolites – in much higher doses than other solid organ transplant recipients, and yet also have the highest rates of acute and chronic allograft rejection. With respect to T cells, these drugs act by inhibiting cell proliferation. In this context, we speculate that CD4^+^CD57^+^PD1^+^ T cells, which exhibit reduced proliferation, escape this inhibition and can continue to produce cytokines. Further work will be required to understand how these cells arise within the lung allograft, and whether specific targeting of these cells may have a therapeutic benefit.

## Methods

### Study design and BAL collection

The samples analyzed in this study were collected with informed consent for research use and were approved by University Health Network Research Ethics Board (15-9531). BAL cells from 50 TLTP lung transplant recipients presenting for surveillance bronchoscopy at 3 months post-transplant were obtained according to a standard protocol ^52^. In addition, for each LT recipient, two further samples were obtained from each recipient to form a longitudinal cohort **(Supp Fig 1**). Cells were centrifuged at 400g for 5 min and resuspended in 1 ml of 10% dimethyl sulfoxide/90% FBS for viably cryopreservation. Cryovials were placed in Mr. Frosty containers at -80°C for 24Lh, then stored in liquid nitrogen until thawing for processing and mass cytometry. BAL cells from a further 14 LT recipients with acute lung allograft dysfunction (ALAD, defined as a decrease in the forced expiratory volume in one second [FEV1] by 10% or more from the baseline FEV1 measurements) were used for sorting fresh CD4^+^CD57^+^PD1^+^ T cells and their CD57^-^PD1^-^ counterparts for functional analyses.

### Mass cytometry

Metal-conjugated antibodies were purchased either preconjugated from Fluidigm or purified from BioLegend, Thermo Fisher Scientific or BD Biosciences and subsequently conjugated to metals using Maxpar Antibody Labeling Kits (Fluidigm) according to the manufacturer’s instructions **(Supp Table 1)**. Cryopreserved BAL cells were recovered and divided into stimulated (with PMA 50ng/mL and ionomycin 500ng/mL) and unstimulated conditions to allow analysis of cytokine secretion expression and surface markers. Cells were incubated with a cocktail of antibodies against cell surface molecules for 30 min at room temperature. After treatment with intracellular fixation and permeabilization buffers (BD Biosciences), cells were incubated with metal-conjugated antibodies against intracellular proteins. Cells were then washed and stained with Intercalator-Ir (Fluidigm) diluted in PBS containing 1.6% paraformaldehyde and stored at 4 °C until next-day acquisition. To assess variation between mass cytometry runs, a control healthy volunteer PBMC sample was aliquoted, recovered and stained and analyzed alongside the BAL samples.

### Mass cytometry data analysis

Mass cytometry data were initially analyzed with manual gating to exclude debris, dead cells and doublets and to select CD45^+^ cells using Cytobank software (Beckman Coulter). We then used the FlowSOM and viSNE algorithms (5,000 iterations, perplexityL=L100) for visualization of high-dimensional data ^21,53^. Subsequently, we applied cluster identification, characterization, and regression (CITRUS) ^22^ on the CD3+ population from the discovery cohort (n=25, **Supp Table 2**) to identify cell subsets associated with ALAD within 30 days of the sample. CITRUS was run at least 5 times with “equal number” of cells per sample, looking for “abundance” as cluster characterization, with a 1% false discovery rate. To incorporate the time-course design of our study, we implemented a generalized linear mixed model analysis-adapted R pipeline as described by Nowicka and colleagues ^23^. In this approach, different timepoints for each LT recipient were defined as a variable to allow intra-subject pairing. Finally, we used manual gating to confirm the results of CITRUS and generalized linear mixed model analysis.

### Imaging Mass Cytometry

Lung tissue samples were obtained during re-transplantation procedures from 8 patients who were undergoing re-transplantation due to end-stage CLAD. A single biopsy was obtained from the upper lobe of each explanted sample. The harvested lung tissue was inflated and immersed in a 10% formalin solution for 24 hours prior to immersion in a 70% ethanol solution for an additional 24-hour period. Samples were then embedded in paraffin. Four-micron thick sections of paraffin-embedded lung tissue were prepared and affixed to slides for further analysis. For every specimen, we identified three distinct 1x1mm Regions of Interest (ROIs) situated around individual airways. In addition to the CLAD patient samples, we included control samples in our study. Controls included lung tissue samples from two lobectomy procedures for suspected cancer and excess tissue from two donor lungs collected at the end of the cold ischemic phase of preservation. Sections of CLAD and control lung tissue were stained with a panel of 35 heavy-metal-tagged antibodies **(Supp Table 6)**. Subsequently, the three ROIs chosen for each specimen were subjected to ablation using the Hyperion Imaging System (Fluidigm) using a laser frequency of 200Hz, with laser power ranging from 3.5 to 4.5. The ablation procedure generated files in both .txt and .mcd formats, which were subsequently subjected to thorough analysis using the HistoCAT software platform^54^.

### Animal model

Orthotopic lung transplantation was performed using C57BL/10 donors and C57BL/6 recipients under different graft preservation conditions as described previously ^27^. Briefly, harvested lungs were stored either under minimal (60 minutes at 4°C followed by 15 min of warm anastomotic time) or prolonged (6h at 4°C, 45 min at 37°C, followed by 15 min of warm anastomotic time) storage prior to reperfusion. Orthotopic lung transplant recipients were sacrificed 28 days after LT for histological analysis and cellular phenotyping. Grafts were flushed with saline and digested with collagenase A and DNase I in a gentleMACS tissue dissociator (Miltenyi Biotec). Cells were stained with cocktails of fluorochrome-conjugated monoclonal antibodies including AF647- conjugated anti-mouse CD57 (polyclonal; Bioss) and AF488-conjugated anti-mouse PD1 (polyclonal; Bioss). Flow cytometry data were analyzed using FlowJo (v10, FlowJo LLC) and correlated with histological grading. We graded peribronchial fibrosis and parenchymal fibrosis in a blinded fashion according to a published scoring system^28^. Immunofluorescent staining of sections of formalin-fixed and paraffin-embedded lung tissue was conducted by antigen retrieval using 1x pH 9 Tris/EDTA buffer before staining with unconjugated anti-mouse CD4. This was followed by a secondary antibody cocktail including Alexa Fluor 555 anti-rabbit IgG (A-31572, Invitrogen) as well as AF647-conjugated anti-mouse CD57 (polyclonal; Bioss) and AF488-conjugated anti-mouse PD1 (polyclonal; Bioss). Nuclei were stained with DAPI and immunofluorescence images were acquired using a Zeiss AxioObserver.

### CITE-seq sample processing and acquisition

We identified four LT recipients scheduled for bronchoscopy with ALAD (FEV_1_ decrease by ≥10% from the prior value). Fresh BAL cells were collected by centrifugation at 400g for 5 minutes. T cells were obtained using a CD3^+^ T cell enrichment kit (EasySep™ Human T Cell enrichment, Stemcell Technologies) according to the manufacturer’s instructions. Enriched T cells were resuspended at 1x10^6^ cells/mL in PBS + 0.05% BSA. 50 µL of the cell suspension were stained with BV711 conjugated anti-human CD4, APC conjugated anti-human PD1 and PE-Cy7 anti-human CD57 and analyzed by flow cytometry to confirm ≥ 7.8% CD57^+^PD1^+^ cells in the BAL CD4^+^ T cell compartment. If confirmed, the remaining cells were used for CITE-seq. Briefly, 1x10^6^ cells were resuspended in 45 μL PBS + 0.05% BSA before adding 5 µL of Fc blocking reagent and incubated for 10 minutes at 4°C. TotalSeq™-C antibody cocktails were prepared during the incubation by adding pre-determined concentrations of each antibody (**Supp Table 6**). The resulting cocktail was then added to cells and incubated for 30 minutes at 4°C. After incubation, the cell pellets were resuspended in 1mL PBS + 0.05% BSA and centrifuged for 5 minutes at 400 x g. After a total of 3 washes, the final cell density was adjusted to 1000-1200 cells/µL and subjected to the 10X Genomics Chromium 5’ Immune Profiling v2.0 protocol for library preparation. A total of 2×10^4^ cells from each sample were loaded onto a 10X single cell G chip. After droplet generation, samples were transferred to a pre-chilled 96 well plate (Eppendorf), heat sealed and incubated overnight in a Veriti 96-well thermocycler (Thermo Fisher). The next day, sample cDNA was recovered using Recovery Agent (10X Genomics) and subsequently cleaned up using Silane DynaBeads (Thermo Fisher) mix as outlined by the user guide 5’ Immune Profiling v2.0. Purified cDNA was amplified for 11 cycles before being cleaned up using SPRIselect beads (Beckman Coulter). Samples were diluted 9:1 (elution buffer (Qiagen):cDNA) and run on a Bioanalyzer (Agilent Technologies) to determine cDNA concentration. The library was then sequenced using an Illumina Novaseq 6000 sequencer using the following parameters Read 1: 28b, index 1: 10bp, Index 2: 10bp, Read 2: 90bp.

### CITE-seq analysis

The sequencing output was first demultiplexed and converted into FASTQ files using Cell Ranger ^55^. Reads were aligned to the human reference genome (grch38), and count UMIs in the mRNA libraries, and CITE-seq-Count to count UMIs in the antibody-derived tag (ADT) libraries. The matrix, barcodes, and gene information from Cell Ranger were then used to create gene-cell matrices and a Seurat object ^56^. For each Seurat object, cells with poor quality were excluded based on the following quality control metrics: unique gene counts over 5000 or below 200, UMI counts over 30,000, and mitochondrial gene expression higher than 10%. Each sample was scaled and normalized using Seurat’s “SCTransform” function followed by adding the protein expression levels to the Seurat object and normalization and scaling for ADT assay. Next, CD4^+^ T cells were subsetted based on high CD4 and low CD8 protein expression (ADT-CD8 < 12 & ADT-CD4 > 12). Seurat objects of CD4+ cells from all four BAL samples were merged into a single Seurat object. The SCTransform method was used for normalization, scaling, and integration which corrects for unwanted sources of variation and enhances the signal of genuine biological differences between cells^57^. Then, dimensionality reduction was applied to the integrated Seurat object, allowing visualization and interpretation of the high-dimensional scRNAseq data. This was followed by unsupervised clustering (resolution = 0.5) which led to the identification of 10 clusters. Then cells with higher expression of protein level of CD57 and PD1 were highlighted on tSNE plot.

RNA velocity estimates the future state of individual cells by calculating the ratio of unspliced to spliced mRNA for each gene. The spliced and unspliced counts from the scRNAseq data were utilized to calculate RNA velocity using the velocyto package in Python^58^. Splicing information extracted from velocyto was then added to all the selected cells in the CD4^+^ Seurat object. Subsequently, CD4^+^ T cells underwent pseudotime trajectory analysis using the scVelo pipeline^59^. This analysis inferred the trajectory and the chronological sequence of cell states, highlighting the differentiation paths taken by cells. The pseudotime values were visualized on the tSNE plots, providing insights into the dynamic processes and cellular transitions at play.

### Fluorescence-activated cell sorting

BAL cells collected from 15 ALAD samples and stained with fluorophore-tagged surface antibodies for 30 min at room temperature. The following antibodies were used to stain the cells: BV711-conjugated anti-human CD4 (BioLegend); APC-conjugated anti-human PD1 (BioLegend); PE/Cy7-conjugated anti-human CD57, in addition to biotinylated CD163, CD14, CD8, and CD56 (all from BioLegend). Cells washed with 500 µL of 1% FBS in PBS and incubated with streptavidin-AF488 (1:200, BioLegend) for 20 minutes, before washing and staining with propidium iodide to identify dead cells. Stained cells were subjected to FACS sorting using a Sony SH800 instrument. After exclusion of PI-positive cells and putative doublets based on forward and side scatter analysis, cells in the dump channel (AF488) were eliminated. The remaining cells were gated to select CD4^+^ T cells and sorted for CD57^+^PD1^+^ and CD57^-^PD1^-^ cells.

### Cell culture, multiplex protein analysis, and *in vitro* assays

Sorted cells were labeled with 1μM CFSE in serum free PBS prior to wash and culture with plate bound anti-CD3 (5μM) and soluble anti-CD28 (1μM) antibodies in DMEM medium containing 5.55 mM D-glucose, 4 mM L-glutamine, and 1 mM sodium pyruvate, supplemented with 10% FBS, 50 U/ml penicillin, and 50 mg/ml streptomycin and 500 U/ml human recombinant IL-2 (Proleukin, Chiron). After 18 hours, 100μL supernatant was replaced with fresh media. The collected supernatant was stored at -80°C for later multiplex analysis. Supernatants were thawed and subjected to a custom multiplex bead kit (R&D Systems) to assess the levels of released analytes including IL-21, IL-4, CXCL8, and granzyme A. Samples were prepared in a 1:1 dilution with assay buffer, as suggested by the manufacturer. Diluted samples and standards were run in duplicate. Biomarker concentrations were obtained using a Bio-Plex MAGPIX Multiplex Reader (Bio-Rad). All samples and standards were run in duplicate according to manufacturers’ protocols.

After 5 days, CFSE dilution in the sorted cells – a measure of T cell proliferation – was determined by flow cytometry. Finally, a restimulation-induced cell death assay^60^ was used to assess whether CD57^+^PD1^+^ cells are less susceptible to this mode of cell death than their CD57^-^PD1^-^ counterparts. CD57^+^PD1^+^ and CD57^-^PD1^-^ cells were stimulated in culture for 7 days before washing, and were then restimulated with anti-CD3 mAb (OKT3, 100 ng/ml final) for 24 hours. Cells were then harvested and stained with Annexin V and propidium iodide for evaluation of apoptotic cell death.

### Statistics

All statistical tests were two-sided unless stated otherwise. The Wilcoxon rank-sum test was used to assess statistical differences between two groups. When assessing >2 groups simultaneously, the nonparametric Kruskal–Wallis test was used. A Fisher’s exact test was applied to assess statistical differences between two categorical variables. Receiver Operating Characteristic (ROC) curves were used to assess classification accuracy, which was quantified by Area Under the Curve (AUC). Kaplan– Meier plots and two-sided log-rank tests were used to compare survival probabilities between groups. Cox Proportional Hazards (PH) regression models were used to adjust for potential confounders for time-to-CLAD or time-to-death/re-transplantation analysis. and Cox regression analyses were used to assess covariates with respect to time-to- allograft dysfunction. All statistical analyses were performed using R v.3.5.1+ or Prism 8+ (GraphPad Software).

## Acknowledgments

The authors wish to thank Martin Oberle, Marcelo Cuesta, Iva Avramov and other members of the Toronto Lung Transplant Program Biobank for their assistance with sample procurement and storage. We also thank Dr. Cynthia Guidos and Dr. Dariush Davani for their critical input on the design of mass cytometry experiments. We also wish to thank Dr. Sarah Crome and Sarah Colpitts for their assistance and advice in planning CITE-Seq experiments.

This work was funded by Cystic Fibrosis Foundation grants JUVET18AB0 and JUVET23G0 (to S.J.), and MARTIN18I0 (to T.M. and S.J.) by an International Society for Heart and Lung Transplantation Norman Shumway Career Development Award (to S.J.), by a Canadian Society of Transplantation Research Training Award (to S.M.) and by the University Health Network Foundation.

## Abbreviations

ALAD: Acute lung allograft dysfunction
CLAD: Chronic lung allograft dysfunction
LT: Lung transplantation
BAL: Bronchoalveolar lavage
IMC: Imaging mass cytometry
tSNE: T-distributed stochastic neighbor embedding
CyTOF: Cytometry by Time of Flight
CITE-seq: Cellular indexing of transcriptomes and epitopes
Tph: peripheral helper T
FEV1: Forced expiratory volume in one second
TLTP: Toronto lung transplant program
GLMM: Generalized linear mixed model
ROC: Receiver-operating characteristic
DSA: Donor specific antibodies
HR: Hazard radio
CMV: Cytomegalovirus
ADT: Antibody derived tags
RICD: Restimulation induced cell death

