## Supplemental material for "Emergence of a senescent and inflammatory pulmonary CD4^+^ T cell population prior to lung allograft failure"

#### Supplementary figure legends

**Supp Fig 1. Study flow diagram for the BAL CyTOF cohort**.

**Supp Fig 2. inter- and intra-patient heterogeneity in BAL cell composition by unsupervised multidimensional clustering.** Uniform Manifold Approximation and Projection plots of BALs from all 3 time points from the discovery group (n=25) using FlowSOM demonstrated high inter- and intra-patient heterogeneity in BAL cell composition. All BAL samples from all 25 patients are shown. Each sample is named according to patient number and BAL sample number (D01_1, D01_2, D01_3, etc.).

**Supp Fig 3. Identification of a BAL $CD3^+$ T cell population associated with lung allograft dysfunction using CITRUS. A.** CITRUS (cluster identification, characterization, and regression) analysis of all BALs from n=25 discovery group patients showing cell populations differentiating patients developing ALAD within 30 days from those whose allograft function remained stable. Red circle indicates the statistically significantly differentially represented population. **B.** Mass cytometry histograms of the top differentially represented population, which expresses CD4, CD57, CD3 and PD1. **C.** Box-and-whisker plot showing the abundance of CITRUS cluster 1076 – representing $CD4^+CD57^+PD\text{-}1^+$ T cells – which exhibited a significant difference between ALAD and stable samples.

**Supp Fig 4. Marker expression by 20 FlowSOM T cell clusters.** Average normalized T cell marker expression (columns) presented for each FlowSOM cluster (rows).

**Supp Fig 5. Temporal relationships between the appearance of $CD4^+CD57^+PD1^+$ T cells and acute and chronic lung allograft dysfunction and in all 50 individuals**. Fifty LT recipients were randomly divided into discovery (1-25) and validation (26-50) groups. Time course of each subject is represented graphically as an orange arrow (arbitrary time scale). For each individual LT recipient, the appearance ≥7.8%

$CD57^+PD1^+$ cells in the BAL $CD4^+$ T cell compartment (blue crosses), the first episode of ALAD (blue triangle) and the onset of CLAD (red triangle).

**Supp Fig 6. $CD4^+CD57^+PD1^+$ T cells are associated with an increased risk of CLAD.** Kaplan-Meier curves showing time to CLAD since appearance of ≥7.8% $CD57^+PD\text{-}1^+$ cells in the BAL $CD4^+$ T cell compartment, or last BAL analyzed. Log rank p=0.0071**.**

**Supp Fig 7. Identification of distinct $CD27^+$ and $CD27^-$ subsets of $CD4^+CD57^+PD1^+$ T cells.** Sequencing by cellular indexing of transcriptomes and epitopes (CITE-Seq) was performed on 4 BAL samples from patients with ALAD. **A.** After downsampling and integration, BAL $CD4^+$ T cells ($CD3^+CD4^+CD8^-$ identified by antibody-derived oligonucleotide tag, ADT) were displayed on a tSNE plot revealing 10 distinct cell clusters (left panel). Feature plots showing PD1 and CD57 protein expression (ADT-PD-1, middle panel and ADT-CD57, right panel) on $CD4^+$ T cells as indicated by purple dots inside red circles. Three distinct clusters with detectable CD57 protein expression are identified (1, 2, 4). **B.** Heat map of the top differentially expressed genes in BAL $CD4^+$ T cells, with clusters identified in A highlighted in red boxes. Cluster 1 contains a $CD27^-$ $LAG3^+$ $CXCL13^+$ $TNFRSF18^+$ population (putative follicular helper function). Cluster 2 contains $CD27^-GZMA^+CXCR6^+CCL5^+$ cells (putative mucosa-associated invariant T cells). Cluster 4 contains $CD27^+GZMK^+NKG7^+EOMES^+$ cells (putative cytotoxic $CD4^+$ T cells). **C.** Feature plots of selected genes differentially expressed in clusters 1, 2 and 4.

**Supp Fig 8. Single-cell TCR clonotype analysis reveals extensive clonal similarity between clusters 1 and 2. A.** Distribution of clonal expansions in BAL $CD4^+$ T cells. Large (20-100) clonal expansions are predominantly seen in clusters 1 and 2. **B.** Analysis of the top 20 expanded clones for each cluster reveals similar proportions of specific clones between clusters 2 and 4; in contrast, few clones are shared with cluster 4, which is composed of mainly distinct clones.

**<u>Supplementary table legends</u>**

**Supp Table 1. Antibody panel for CyTOF.** Rows highlighted in yellow contain intracellular markers.

**Supp Table 2. Clinical characteristics of participants.** Patients were randomized into a discovery group (n=25) and a validation group (n=25). LT, lung transplant; BLT, bilateral lung transplant; CMV, cytomegalovirus; D, donor; R, recipient.

**Supp Table 3. Clinical and bronchoscopic parameters associated with ≥ 7.8% $CD57^+PD1^+$ T cells in the BAL $CD4^+$ T cell compartment.** Fisher's exact tests were performed. [1]Missing samples excluded. [2]Comparison of normal to abnormal cytology. [3]Comparison of A0/AX with A1 or greater. [4]Testing not done (n.d.) due to rarity of small airway inflammation in the cohort. [5]Comparison of any antimicrobial therapy pre- or post-bronchoscopy vs. none. [6]Testing not done (n.d.) due to small number of events. *Eosinophilia noted in one of the samples with lymphocytosis. **Eosinophilia noted in isolation in one case and in another sample with neutrophilia. †One patient received both pre-bronchoscopy antibiotics and post-bronchoscopy antivirals.

**Supp Table 4. Change in immunosuppression in patients with and without ≥ 7.8% $CD57^+PD1^+$ T cells in the $CD4^+$ T cell compartment of at least one BAL sample.** Fisher's exact tests were performed. CNI, calcineurin inhibitor; ATG, anti-thymocyte globulin.

**Supp Table 5. Comparison of clinical characteristics and survival of patients with and without ≥ 7.8% $CD57^+PD1^+$ cells in the BAL $CD4^+$ T cell compartment. A.** Clinical characteristics of patients with and without ≥ 7.8% $CD57^+PD\text{-}1^+$ cells, excluding 2 patients who developed CLAD prior to the appearance of the cells. **B.** Univariable and bivariable time-dependent Cox proportional hazards models for time to CLAD. **C.** Univariable and bivariable time-dependent Cox proportional hazards models for time to death or re-transplantation.

**Supp Table 6. Antibody panel for imaging mass cytometry.** Poly, polyclonal.

**Supp Table 7. Oligonucleotide-conjugated antibody panel for CITE-seq.**

**Supplementary Figure 1**

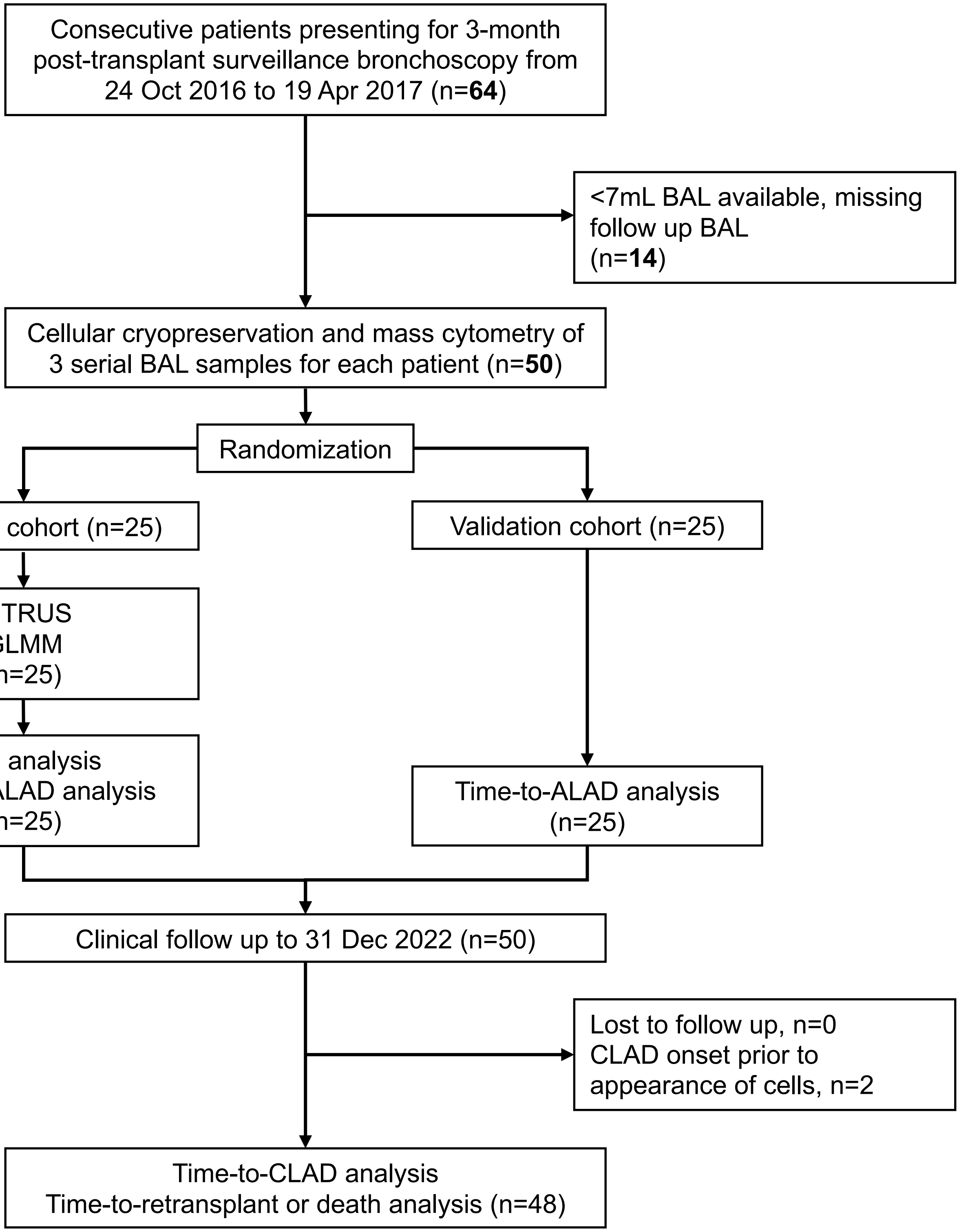

### Supplementary Figure 2

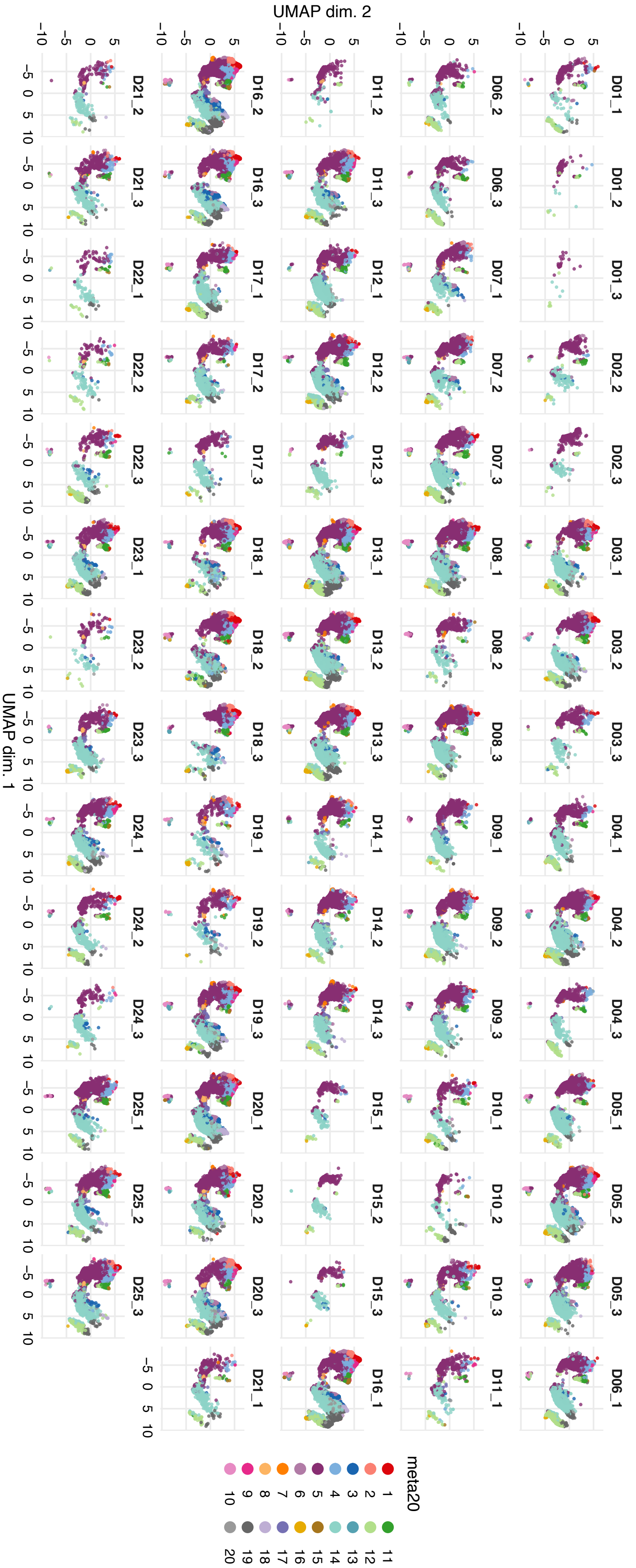

Supplementary Figure 3

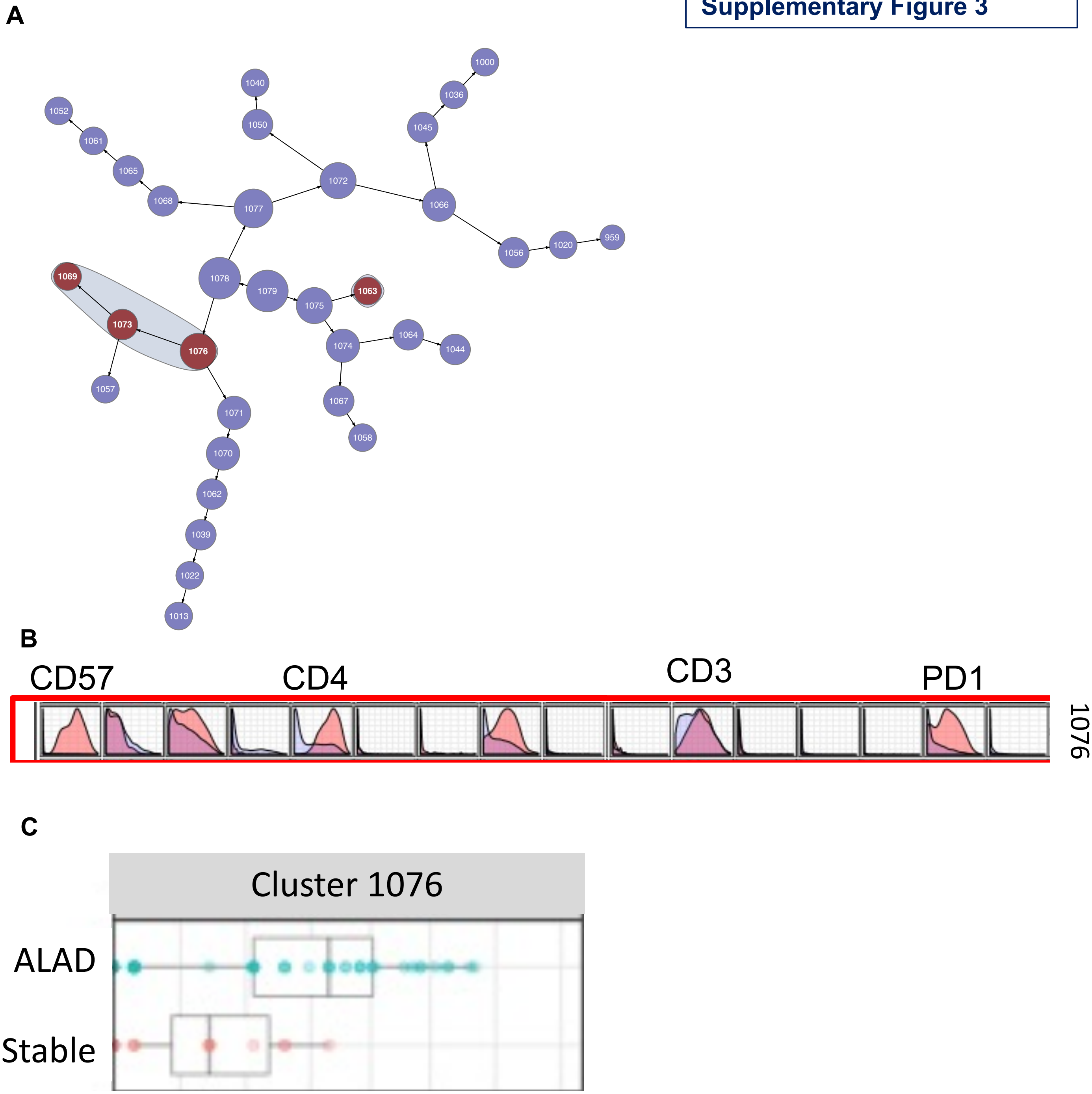

#### Supplementary Figure 4

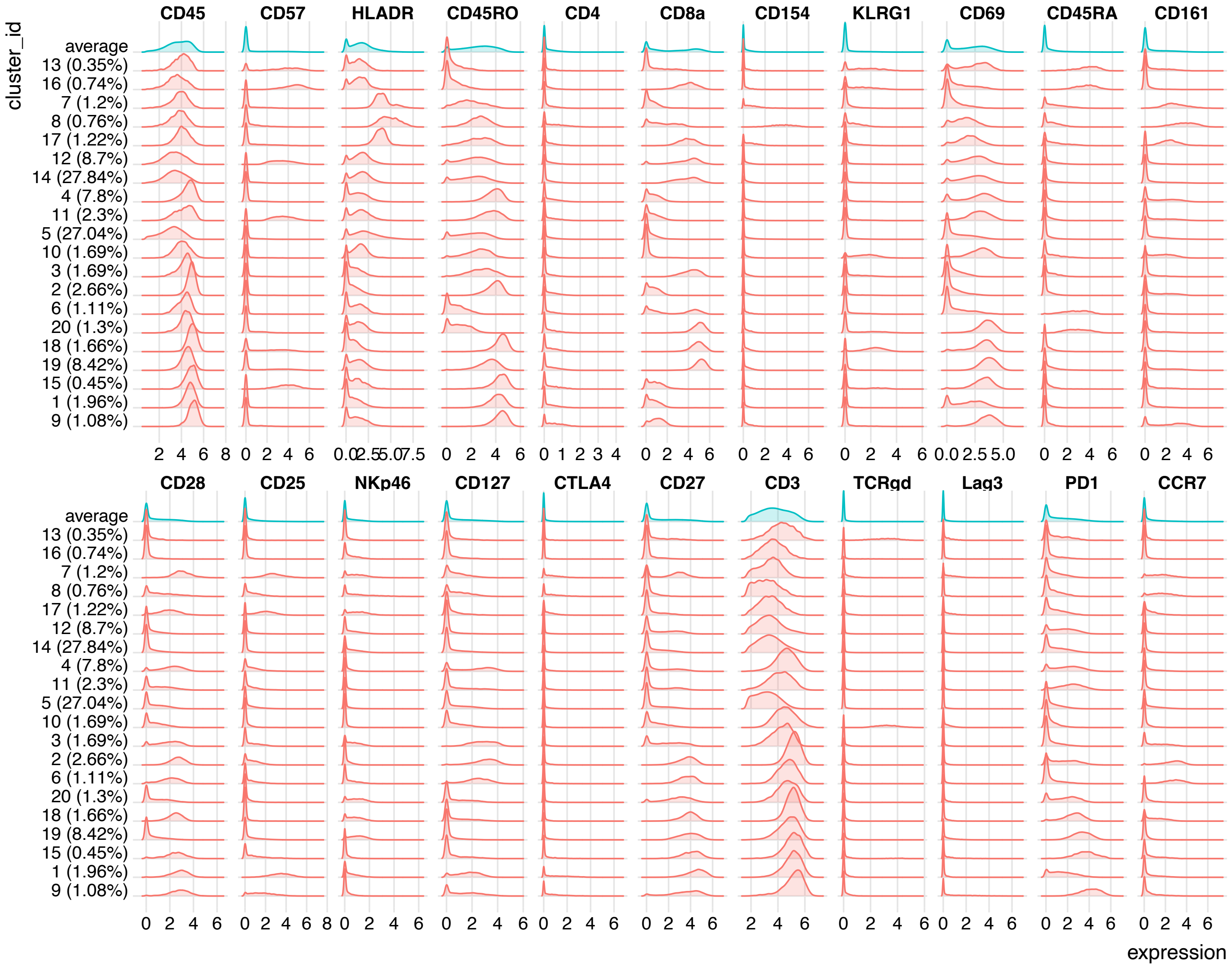

**Supplementary Figure 5**

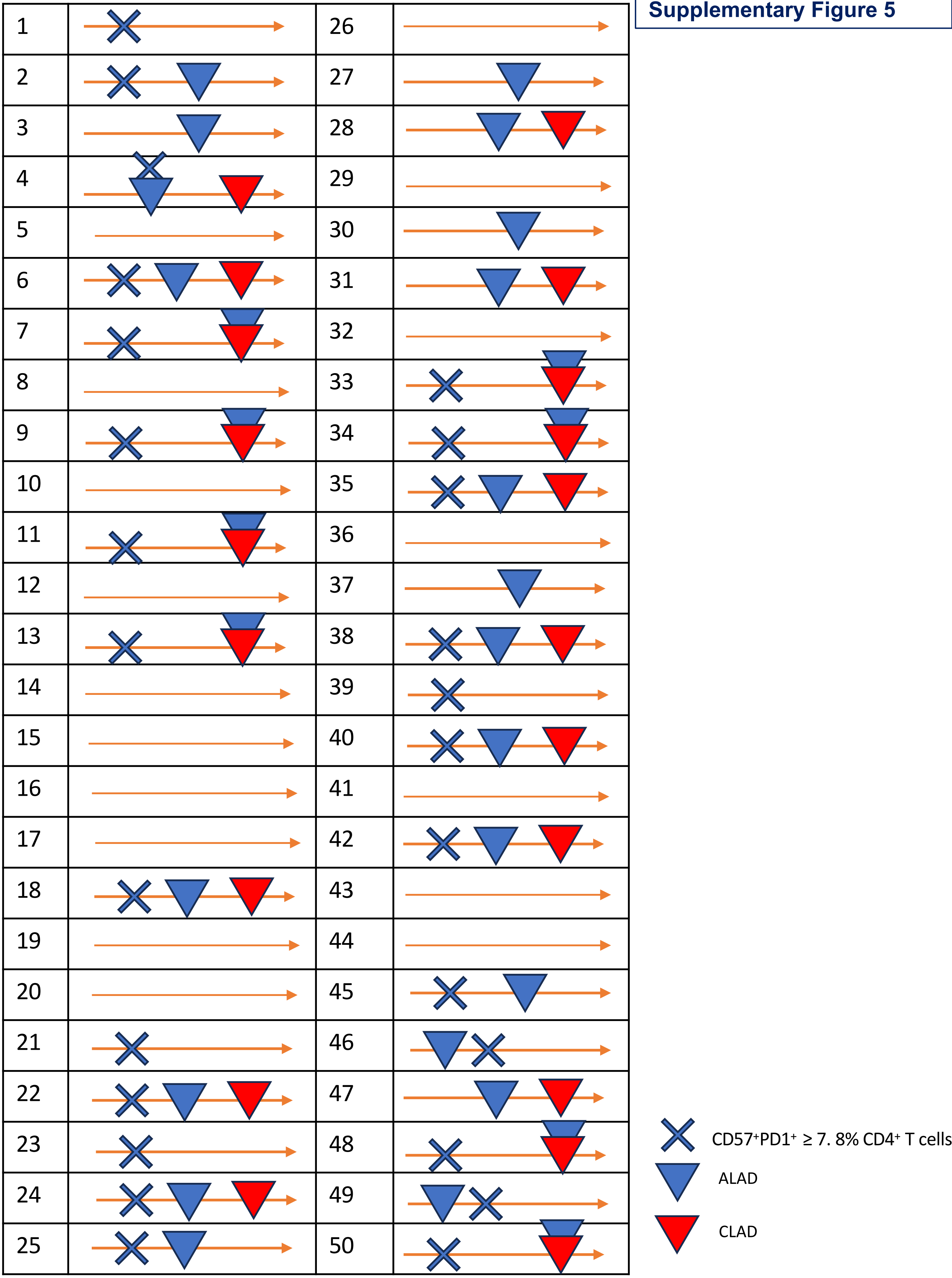

**Supplementary Figure 6**

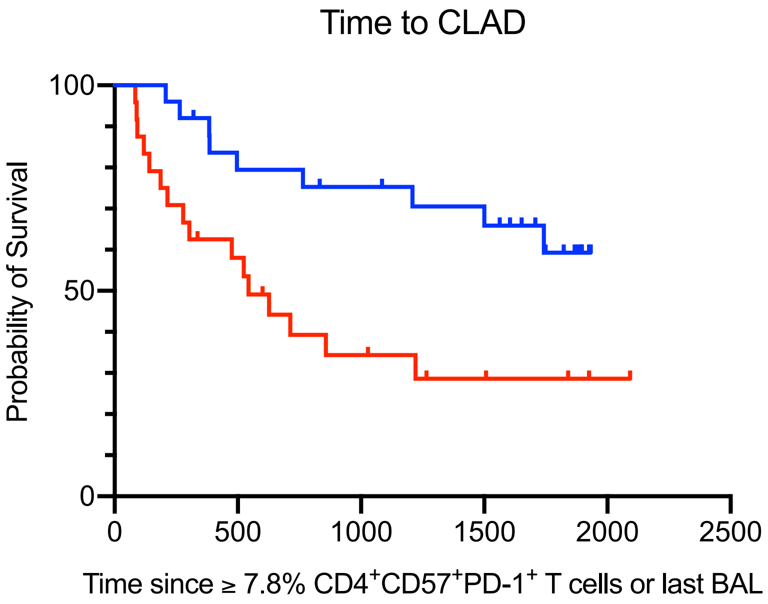

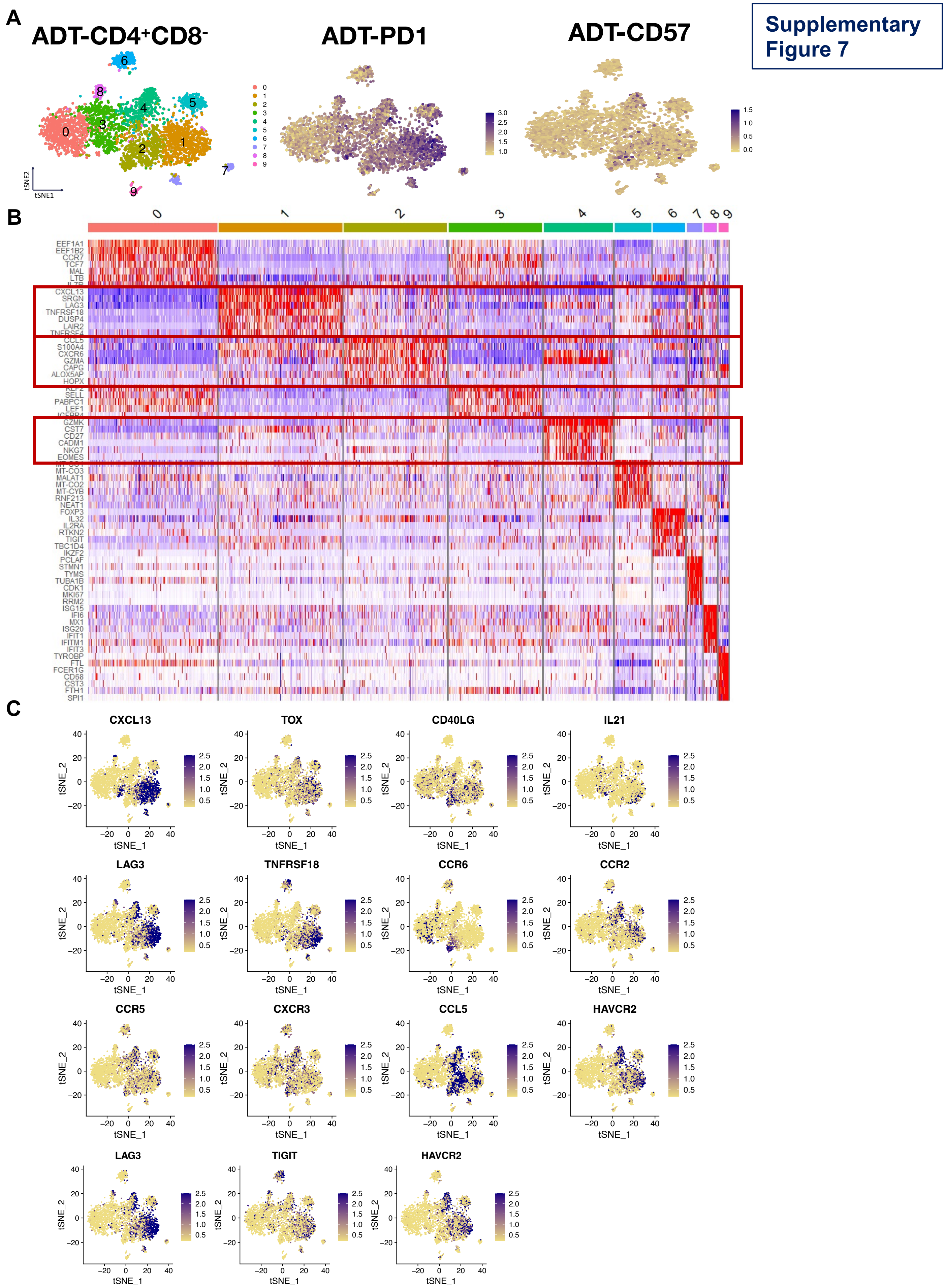

Supplementary Figure 7
A
ADT-CD4+CD8-
ADT-PD1
ADT-CD57
tSNE1
tSNE2
B
EEF1A1
EEF1B2
CCR7
TCF7
MAL
LTB
CXCL13
SRGN
LAG3
TNFRSF18
DUSP4
LAIR2
S100A4
CXCR6
GZMA
CAPG
ALOX5AP
HOPX
SELL
PABPC1
LEF1
GZMK
CST7
CD27
CADM1
NKG7
EOMES
MT-CO3
MALAT1
MT-CO2
MT-CYB
RNF213
NEAT1
FOXP3
IL32
IL2RA
RTKN2
TIGIT
TBC1D4
IKZF2
PCLAF
STMN1
TYMS
TUBA1B
CDK1
MKI67
RRM2
ISG15
IFI6
MX1
ISG20
IFIT1
IFITM1
IFIT3
TYROBP
FTL
FCER1G
CD68
CST3
FTH1
SPI1
C
CXCL13
TOX
CD40LG
IL21
LAG3
TNFRSF18
CCR6
CCR2
CCR5
CXCR3
CCL5
HAVCR2
LAG3
TIGIT
HAVCR2
tSNE_1
tSNE_2

**Supplementary Figure 8**

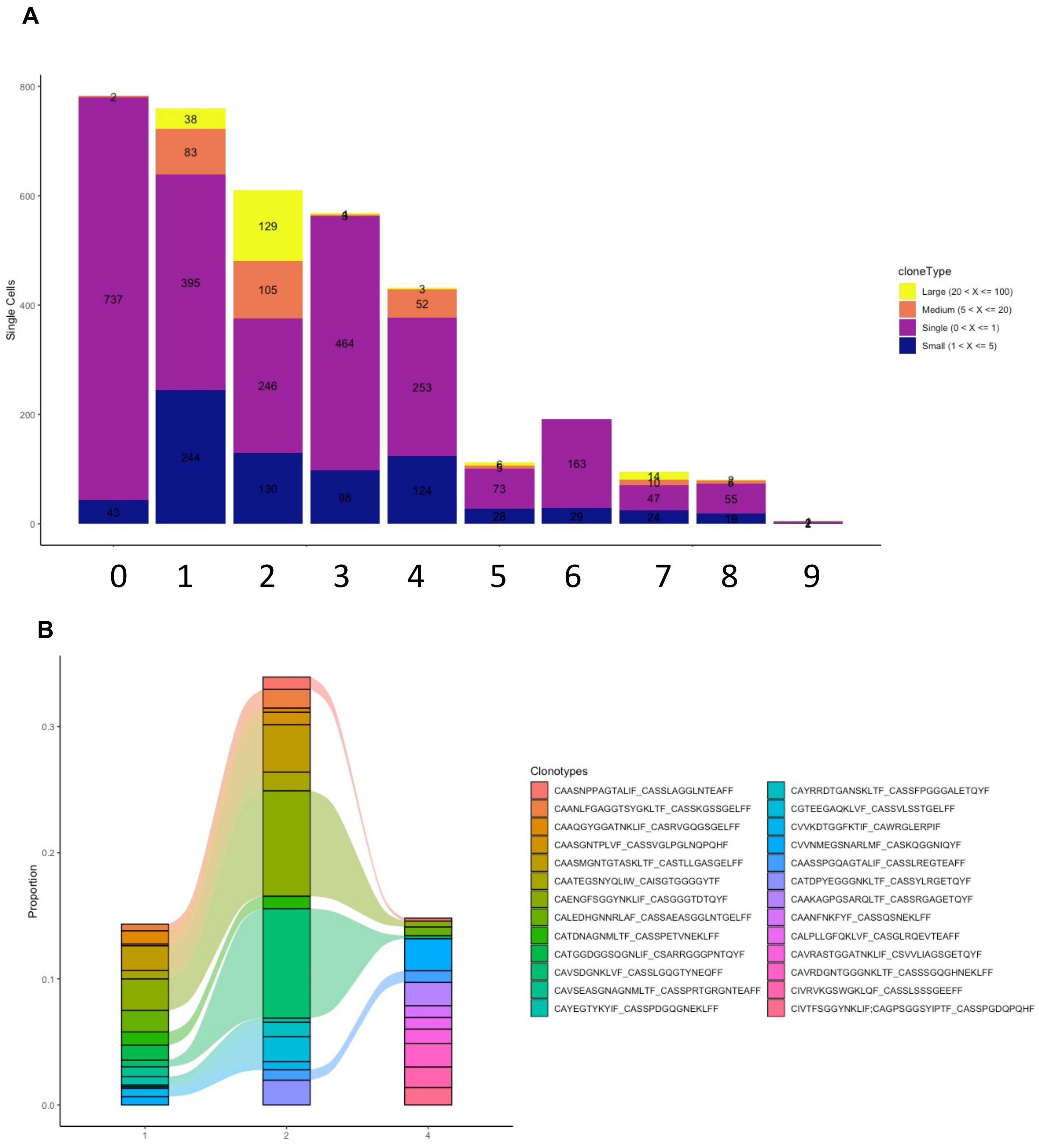

#### Supplementary Table 1

| Tag | Specificity | Clone |
|---|---|---|
| 89Y | CD45 | HI30 |
| 115In | CD57 | HCD57 |
| 141Pr | HLA-DR | L243 |
| 142Nd | CD45-RO | UCHL1 |
| 143Nd | CD11c | Bu15 |
| 145Nd | CD4 | SK3 |
| 146Nd | CD8a | RPA-T8 |
| 147Sm | Epcam | 9C4 |
| 148Nd | CD154 | 24-31 |
| 149Sm | KLRG1 | SA231A2 |
| 150Nd | CD69 | FN50 |
| 151Eu | CD123 | 6H6 |
| 152Sm | TNFα | Mab11 |
| 153Eu | CD45RA | HI100 |
| 154Sm | CD19 | HIB19 |
| 155Gd | CD161 | HP-3G10 |
| 156Gd | CD28 | CD28.2 |
| 158Gd | CD33 | WM53 |
| 159Tb | CD90 | 5 E 10 |
| 160Gd | CD14 | M5E2 |
| 161Dy | CD25 | M-A251 |
| 162Dy | NKp46 | 9E2 |
| 163Dy | CD127 | eBioRDR5 |
| 164Dy | IL17A | BL168 |
| 165Ho | CTLA-4 | |
| 166Er | Siglec 8 | 7C9 |
| 167Er | CD27 | O323 |
| 168Er | IFNγ | B27 |
| 169Tm | IL10 | JES3-19F1 |
| 170Er | CD3 | UCHT1 |
| 171Yb | CD163 | GHI/61 |
| 172Yb | TCRgd | 5A6.E9 |
| 173Yb | Lag3 | 11C3C65 |
| 174Yb | CD56 | NCAM16.2 |
| 175Lu | CD279, PD1 | EH12.2H7 |
| 176Yb | CCR7 | G043H7 |
| 209Bi | CD16 | |

#### Supplementary Table 2

| | Discovery (n=25) | Validation (n=25) | P value |
|---|---|---|---|
| Recipient age at LT, years | 61 | 58 | 0.95 |
| Male Sex | 17 | 18 | 0.96 |
| Donor-recipient sex mismatch | 2 | 4 | 0.84 |
| Type of TX (BLT) | 19 | 19 | 1.0 |
| Native lung disease | | | 0.57 |
| Pulmonary fibrosis | 11 | 12 | |
| Chronic obstructive pulmonary disease | 9 | 6 | |
| Other | 5 | 7 | |
| CMV Serology | | | 1.0 |
| D+/R- | 4 | 5 | |
| D+/R+, D-/R+, D-/R- | 21 | 20 | |

#### Supplementary Table 3

| Features Present at Each Bronchoscopy | %CD57+PD-1+ in BAL CD4+ T cell compartment | | | | | | | | |
|---|---|---|---|---|---|---|---|---|---|
| | First BAL | | | Second BAL | | | Third BAL | | |
| | <7.8% (n=34) | ≥7.8% (n=16) | p | <7.8% (n=42) | ≥7.8% (n=8) | p | <7.8% (n=31) | ≥7.8% (n=19) | p |
| Clinical symptoms | | | | | | | | | |
| Absent | 23 | 13 | 0.50 | 24 | 3 | 0.44 | 16 | 10 | >0.99 |
| Present | 11 | 3 | | 18 | 5 | | 15 | 9 | |
| Respiratory | 9 | 2 | | 15 | 5 | | 11 | 9 | |
| Systemic | 1 | 1 | | 5 | 1 | | 4 | 4 | |
| Upper GI | 2 | 1 | | 3 | 0 | | 2 | 1 | |
| BAL microbiology | | | | | | | | | |
| Category 1,2 | 6 | 4 | 0.71[1] | 5 | 1 | >0.99[1] | 4 | 1 | 0.63[1] |
| Category 3,4,5 | 26 | 12 | | 35 | 7 | | 27 | 18 | |
| Missing | 2 | 0 | | 2 | 0 | | 0 | 0 | |
| BAL cytology | | | | | | | | | |
| Negative | 21 | 10 | >0.99[2] | 31 | 4 | 0.22[2] | 17 | 10 | >0.99[2] |
| Neutrophilia | 8 | 5 | | 5 | 1 | | 6 | 5 | |
| Lymphocytosis | 3 | 0 | | 2 | 1 | | 6 | 2 | |
| Mixed Neut/Lymph | 2 | 1 | | 3 | 2 | | 2 | 1 | |
| Eosinophilia | 0 | 0 | | 1 | 0 | | 1* | 2** | |
| ISHLT A grade | | | | | | | | | |
| A0 | 23 | 8 | 0.68[3] | 24 | 4 | >0.99[3] | 16 | 9 | >0.99[3] |
| AX | 5 | 3 | | 12 | 3 | | 7 | 6 | |
| A1 | 5 | 2 | | 3 | 1 | | 1 | 2 | |
| ≥A2 | 0 | 1 | | 2 | 0 | | 1 | 0 | |
| Biopsy not performed | 1 | 2 | | 1 | 0 | | 6 | 2 | |
| ISHLT B grade | | | | | | | | | |
| B0 | 13 | 1 | n.d.[4] | 15 | 3 | n.d.[4] | 11 | 6 | n.d.[4] |
| BX | 20 | 13 | | 26 | 5 | | 14 | 10 | |
| B1R | 0 | 0 | | 0 | 0 | | 0 | 1 | |
| B2R | 0 | 0 | | 0 | 0 | | 0 | 0 | |
| Biopsy not performed | 1 | 2 | | 1 | 0 | | 6 | 2 | |
| Antimicrobial therapy | | | | | | | | | |
| None | 18 | 6 | 0.38[5] | 28 | 5 | >0.99[5] | 23 | 8 | 0.04[5] |
| Post-bronchoscopy | 5 | 1 | | 7 | 0 | | 5 | 2 | |
| Antibiotics | 4 | 1 | | 4 | 0 | | 2 | 0 | |
| Antifungals | 0 | 0 | | 2 | 0 | | 2 | 1 | |
| Antivirals | 1 | 0 | | 1 | 0 | | 1 | 1 | |
| Pre-bronchoscopy | 12† | 9 | | 7 | 3 | | 4 | 9 | |
| Altered immunosuppression within 30 days of bronchoscopy | | | | | | | | | |
| CNI switch | 2 | 3 | n.d.[6] | 2 | 1 | n.d.[6] | 1 | 1 | n.d.[6] |
| Corticosteroid bolus | 0 | 1 | | 1 | 1 | | 2 | 0 | |
| Anti-thymocyte globulin | 0 | 0 | | 0 | 0 | | 1 | 1 | |

### Supplementary Table 4

| Change in Immunosuppression | %CD57+PD-1+ in BAL CD4+ T cell compartment | | |
|---|---|---|---|
| | < 7.8% in all samples | ≥ 7.8% in at least one sample | p |
| CNI switch | 3 | 7 | 0.29 |
| Augmentation (steroid bolus or ATG) | 3 | 4 | >0.99 |

### Supplementary Table 5

**A**

| | No exposure | Exposure | p |
|---|---|---|---|
| n | 24 | 24 | |
| Age.at.transplant (median [IQR]) | 56.72 [52.23, 64.94] | 60.55 [49.04, 66.60] | 0.934 |
| Sex = M (%) | 18 (75.0) | 16 (66.7) | 0.751 |
| CMV.MISMATCH = 1 (%) | 8 (33.3) | 6 (25.0) | 0.751 |
| Primary.Disease.Category (%) | | | 0.572 |
| CF/bronchiectasis | 2 ( 8.3) | 4 (16.7) | |
| Fibrosis/restrictive | 15 (62.5) | 12 (50.0) | |
| Obstructive | 7 (29.2) | 7 (29.2) | |
| Other | 0 ( 0.0) | 1 ( 4.2) | |
| clad = 1 (%) | 9 (37.5) | 16 (66.7) | 0.083 |
| deathReTx = 1 (%) | 5 (20.8) | 17 (70.8) | 0.001 |

**B**

Univariable time-dependent Cox model for Time to CLAD

| | HR | 2.5 % | 97.5 % | p_value |
|---|---|---|---|---|
| exposure1 | 3.611 | 1.554 | 8.392 | <0.01 |

Bivariable time-dependent Cox model for Time to CLAD

| | HR | 2.5 % | 97.5 % | p_value |
|---|---|---|---|---|
| exposure1 | 3.665 | 1.572 | 8.545 | <0.01 |
| Age.at.transplant | 0.991 | 0.963 | 1.021 | 0.563 |

Bivariable time-dependent Cox model for Time to CLAD

| | HR | 2.5 % | 97.5 % | p_value |
|---|---|---|---|---|
| exposure1 | 3.448 | 1.472 | 8.074 | <0.01 |
| SexM | 0.598 | 0.266 | 1.345 | 0.214 |

Bivariable time-dependent Cox model for Time to CLAD

| | HR | 2.5 % | 97.5 % | p_value |
|---|---|---|---|---|
| exposure1 | 4.021 | 1.699 | 9.516 | <0.01 |
| CMV.MISMATCH1 | 1.89 | 0.821 | 4.351 | 0.135 |

**C**

Univariable time-dependent Cox model for Time to Death/ReTx

| | HR | 2.5 % | 97.5 % | p_value |
|---|---|---|---|---|
| exposure1 | 4.82 | 1.772 | 13.1 | <0.01 |

Bivariable time-dependent Cox model for Time to Death/ReTx

| | HR | 2.5 % | 97.5 % | p_value |
|---|---|---|---|---|
| exposure1 | 4.816 | 1.771 | 13.1 | <0.01 |
| Age.at.transplant | 1.021 | 0.983 | 1.06 | 0.282 |

Bivariable time-dependent Cox model for Time to Death/ReTx

| | HR | 2.5 % | 97.5 % | p_value |
|---|---|---|---|---|
| exposure1 | 4.835 | 1.77 | 13.2 | <0.01 |
| SexM | 1.031 | 0.418 | 2.544 | 0.948 |

Bivariable time-dependent Cox model for Time to Death/ReTx

| | HR | 2.5 % | 97.5 % | p_value |
|---|---|---|---|---|
| exposure1 | 4.927 | 1.806 | 13.4 | <0.01 |
| CMV.MISMATCH1 | 1.314 | 0.532 | 3.246 | 0.553 |

#### Supplementary Table 6

| Tag | Marker | Clone |
|---|---|---|
| 141Pr | **alphaSMA** | 1A4 |
| 142Nd | **CD68** | KP1 |
| 143Nd | **Vimentin** | RV202 |
| 144Nd | **CD14** | EPR3653 |
| 145Nd | **CD31** | EPR3094 |
| 146Nd | **CD16** | EPR16784 |
| 147Sm | **CD163** | EDHu-1 |
| 148Nd | **Pan-keratin** | C11 |
| 149Sm | **CD11b** | EPR1344 |
| 150Nd | **PD-L1** | E1L3N |
| 151Eu | **IgM** | Poly Rb |
| 152Sm | **CD45** | D9M8I |
| 153Eu | **CD138** | SDC1 |
| 154Sm | **CD11c** | Poly |
| 155Gd | **FoxP3** | 236A/E7 |
| 156Gd | **CD4** | EPR6855 |
| 158Gd | **p-STAT3** | 4/P-STAT3 |
| 159Tb | **p21** | 12D1 |
| 160Gd | **Vista** | D1L2G |
| 161Dy | **CD20** | H1 |
| 162Dy | **CD8a** | C8/144B |
| 163Dy | **CD196/CCR6** | G034E3 |
| 164Dy | **CCSP** | 394324 |
| 165Ho | **PD-1** | EPR4877(2) |
| 166Er | **CD141** | M80 |
| 167Er | **GranzymeB** | EPR20129-217 |
| 168Er | **Ki-67** | B56 |
| 169Tm | **Collagen I** | Goat Poly |
| 170Er | **CD3** | Poly |
| 171Yb | **MPO** | E1E71 |
| 172Yb | **CD57** | HCD57 |
| 173Yb | **CD45RO** | UCHL1 |
| 174Yb | **E-cadherin** | 24E 10 |
| 175Lu | **CD25** | EPR6452 |
| 176Yb | **p40** | Poly |
| 191/3Ir | **Intercalator** | |

#### Supplementary Table 7

| Antibody ID | Description | Clone | Barcode | volume per 1 million (ul) |
|---|---|---|---|---|
| TotalSeq™-C0034 | anti-humanCD3 | UCHT1 | CTCATTGTAACTCCT | 0.025 |
| TotalSeq™-C0063 | anti-humanCD45RA | HI100 | TCAATCCTTCCGCTT | 0.0625 |
| TotalSeq™-C0072 | anti-humanCD4 | RPA-T4 | TGTTCCCGCTCAACT | 0.05 |
| TotalSeq™-C0080 | anti-humanCD8a | RPA-T8 | GCTGCGCTTTCCATT | 0.05 |
| TotalSeq™-C0085 | anti-humanCD25 | BC96 | TTTGTCCTGTACGCC | 0.05 |
| TotalSeq™-C0087 | anti-humanCD45RO | UCHL1 | CTCCGAATCATGTTG | 0.25 |
| TotalSeq™-C0088 | anti-humanCD279 (PD-1) | EH12.2H7 | ACAGCGCCGTATTTA | 0.25 |
| TotalSeq™-C0145 | anti-humanCD103 (IntegrinαE) | Ber-ACT8 | GACCTCATTGTGAAT | 1 |
| TotalSeq™-C0146 | anti-humanCD69 | FN50 | GTCTCTTGGCTTAAA | 0.05 |
| TotalSeq™-C0148 | anti-humanCD197 (CCR7) | G043H7 | AGTTCAGTCAACCGA | 1 |
| TotalSeq™-C0154 | anti-humanCD27 | O323 | GCACTCCTGCATGTA | 0.085 |
| TotalSeq™-C0159 | anti-humanHLA-DR | L243 | AATAGCGAGCAAGTA | 0.212 |
| TotalSeq™-C0168 | anti-humanCD57 | QA17A04 | AACTCCCTATGGAGG | 0.05 |
| TotalSeq™-C0386 | anti-humanCD28 | CD28.2 | TGAGAACGACCCTAA | 0.0625 |
| TotalSeq™-C0390 | anti-humanCD127 (IL-7Rα) | A019D5 | GTGTGTTGTCCTATG | 0.05 |
